# How spiders make their eyes: Systemic paralogy and function of retinal determination network homologs in arachnids

**DOI:** 10.1101/2020.04.28.067199

**Authors:** Guilherme Gainett, Jesús A. Ballesteros, Charlotte R. Kanzler, Jakob T. Zehms, John M. Zern, Shlomi Aharon, Efrat Gavish-Regev, Prashant P. Sharma

## Abstract

Arachnids are important components of cave ecosystems and display many examples of troglomorphisms, such as blindness, depigmentation, and elongate appendages. Little is known about how the eyes of arachnids are specified genetically, let alone the mechanisms for eye reduction and loss in troglomorphic arachnids. Additionally, paralogy of Retinal Determination Gene Network (RDGN) homologs in spiders has convoluted functional inferences extrapolated from single-copy homologs in pancrustacean models. Here, we investigated a sister species pair of Israeli cave whip spiders (Arachnopulmonata, Amblypygi, *Charinus*) of which one species has reduced eyes. We generated the first embryonic transcriptomes for Amblypygi, and discovered that several RDGN homologs exhibit duplications. We show that paralogy of RDGN homologs is systemic across arachnopulmonates (arachnid orders that bear book lungs), rather than being a spider-specific phenomenon. A differential gene expression (DGE) analysis comparing the expression of RDGN genes in field-collected embryos of both species identified candidate RDGN genes involved in the formation and reduction of eyes in whip spiders. To ground bioinformatic inference of expression patterns with functional experiments, we interrogated the function of three candidate RDGN genes identified from DGE in a spider, using RNAi in the spider *Parasteatoda tepidariorum.* We provide functional evidence that one of these paralogs, *sine oculis/Six1 A* (*soA*), is necessary for the development of all arachnid eye types. Our results support the conservation of at least one RDGN component across Arthropoda and establish a framework for investigating the role of gene duplications in arachnid eye diversity.

## Introduction

Cave habitats offer apt systems for investigating the genetic basis of morphological convergence because communities of these habitats are similarly shaped by environmental pressures, such as absence of light and diminished primary productivity (Howarth, 1993; Juan, Guzik, Jaume, & Cooper, 2010). Troglobites, species exclusive to cave environments and adapted to life in the dark, exhibit a suite of characteristics common to cave systems around the world, such as reduction or complete loss of eyes, depigmentation, elongation of appendages and sensory structures, and decreased metabolic activity (Jemec, Škufca, Prevorčnik, Fišer, & Zidar, 2017; Protas & Jeffery, 2012; Riddle et al., 2018). Previous work has shown that troglomorphism can evolve over short time spans (<50 kyr) despite gene flow (Bradic, Teotónio, & Borowsky, 2013; Coghill, Darrin Hulsey, Chaves-Campos, García de Leon, & Johnson, 2014; Herman et al., 2018) and that parallel evolution of troglomorphic traits (e.g., depigmentation; eye loss) in independent populations can involve the same genetic locus (Protas et al., 2005; Protas, Trontelj, & Patel, 2011; Re et al., 2018).

Troglomorphism and troglobitic fauna have been analyzed across numerous taxonomic groups with respect to systematics and population genetics. However, one component of the troglobitic fauna that remains poorly understood is cave arachnids. Most orders of Arachnida are prone to nocturnal life history and some orders broadly exhibit troglophily; in fact, troglobitic species are known from all the extant terrestrial arachnid orders except Solifugae and Uropygi (Cruz-López, Proud, & Pérez-González, 2016; Esposito et al., 2015; Harvey, 2002; 2007; Hedin & Thomas, 2010; Mammola, Mazzuca, Pantini, Isaia, & Arnedo, 2017; Miranda, Aharon, Gavish-Regev, Giupponi, & Wizen, 2016; Santibáñez López, Francke, & Prendini, 2014; Smrž, Kováč, Mikeš, & Lukešová, 2013). In addition to eye and pigment loss, troglomorphism in arachnids manifests in the form of compensatory elongation of walking legs and palps, appendages which harbor sensory structures in this group (Derkarabetian, Steinmann, & Hedin, 2010; Mammola & Isaia, 2017; Mammola et al., 2018a; Mammola, Cardoso, Ribera, Pavlek, & Isaia, 2018b).

Thorough understanding of the developmental genetic basis for the evolution of troglomorphic traits has been largely spearheaded in two model systems: the Mexican cave fish *Astyanax mexicanus* (Bradic et al., 2013; Coghill et al., 2014; Herman et al., 2018; Porter, Dittmar, & Pérez-Losada, 2007; Protas et al., 2005; Protas & Jeffery, 2012) and the cave isopod *Asellus aquaticus* (Jemec et al., 2017; Re et al., 2018; Stahl et al., 2015). Both model systems have more than one hypogean population, can be maintained in laboratories, and are amenable to approaches such as genetic crosses and quantitative trait locus mapping. The advent of short read sequencing technology in tandem with experimental approaches has transformed the potential to triangulate regulatory differences between hypogean (subterranean) and epigean (surface-dwelling) lineages (Protas et al., 2005; Re et al., 2018; Riddle et al., 2018; Stahl et al., 2015), and to study a broader range of cave taxa.

Among arthropods, work on the isopod *A. aquaticus* in particular has made significant advances in the identification of loci regulating pigmentation and size of arthropod eyes (Protas et al., 2011; Re et al., 2018), complementing forward and reverse genetic screening approaches in other pancrustacean models (e.g., *Drosophila melanogaster*, *Tribolium castaneum*, and *Gryllus bimaculatus*) (Cagan, 2009; Kumar, 2009; Takagi et al., 2012; ZarinKamar et al., 2011). However, developmental and genetic insights into the evolution of blindness illuminated by *A. aquaticus* and other pancrustacean models are not directly transferable to Arachnida for two reasons. First, the eyes of arachnids are structurally and functionally different from those of pancrustaceans. Typically, the main eyes of adult Pancrustacea (e.g., *A. aquaticus*) are a pair of faceted (or apposition) eyes, which are composed of many subunits of ommatidia. In addition, adult Pancrustacea have small median ocelli (typically three in holometabolous insects), often located medially and at the top of the head.

By contrast, extant arachnids lack ommatidia and typically have multiple pairs of eyes arranged along the frontal carapace. All arachnid eyes are simple-lens eyes or ocelli; each eye has a single cuticular lens, below which are a vitreous body and visual cells. The retina is composed of the visual cells and pigment cells. These eyes are divided in two types, namely the principal eyes and the secondary eyes (Foelix, 2011; Land, 1985). Principal and secondary eyes differ in the orientation of their retina (Homann, 1971): the principal eyes are of the everted type, with the visual cells lying distally, and lack a reflective layer; the secondary eyes are inverted, with the light-sensitive rhabdomeres pointing away from incoming light (analogous to vertebrate eyes). All secondary eyes possess a reflective layer of crystalline deposits called a tapetum, which is responsible for the “eye shine” of spiders. The principal eyes are the median eyes (ME, also known as anterior medium eyes). The secondary eyes comprise the anterior lateral eyes (ALE), posterior lateral eyes (PLE), and medium lateral eyes (MLE; also known as posterior medium eyes) (Fig. 1A) (Foelix, 2011; Land, 1985) (nomenclature used here follows Schomburg et al 2015). Certain orders and suborders of arachnids have lost one type of eye altogether, with the homology of eyes clarified by the fossil record and embryology (Foelix, 2011; Garwood, Sharma, Dunlop, & Giribet, 2014; Morehouse, Buschbeck, Zurek, Steck, & Porter, 2017).

**Figure 1:**
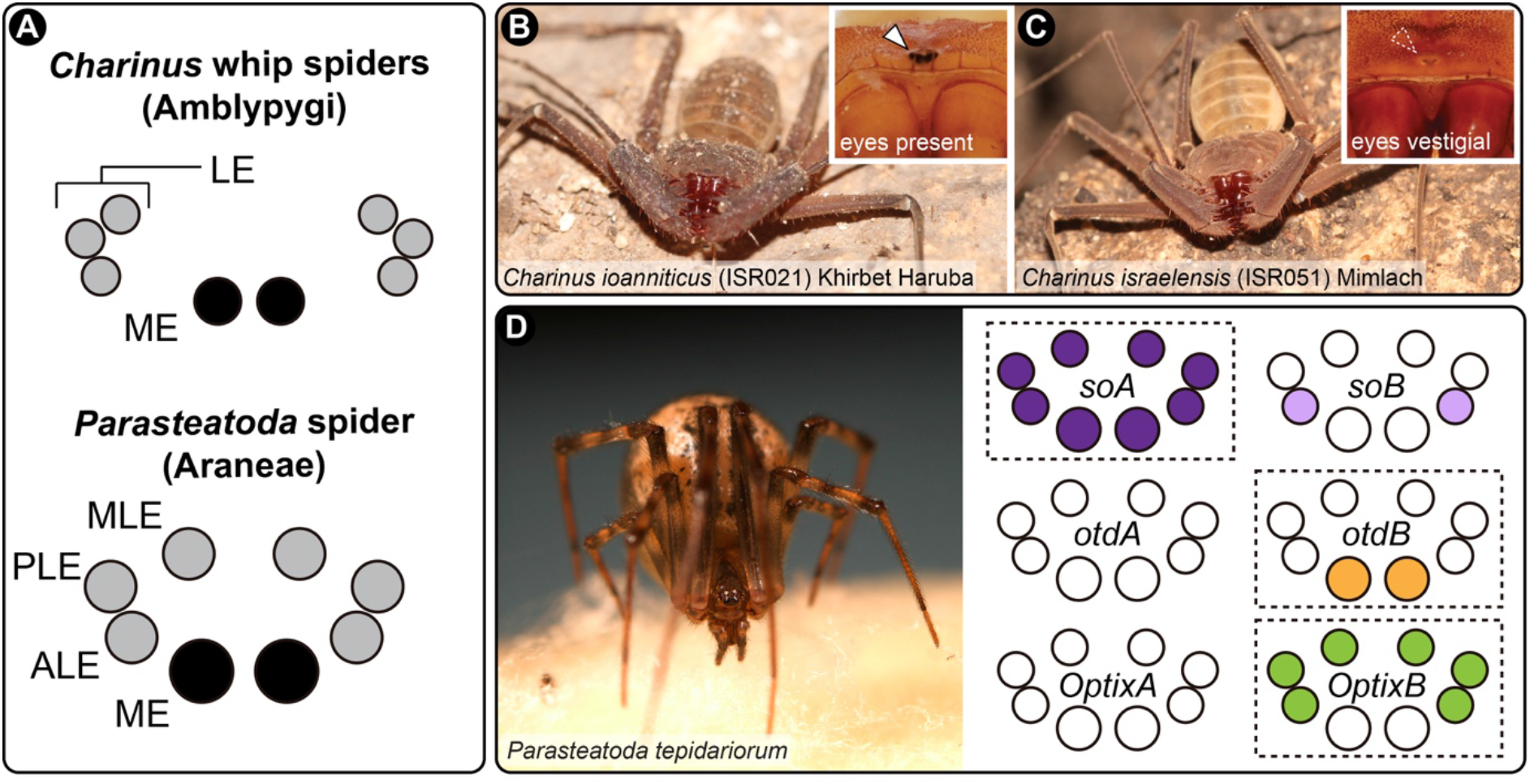
Species used in this study and their eye arrangement. A: Schematic representation of the eyes of *Charinus* whip spiders (Amblypygi) (upper), and the spider *Parasteatoda tepidariorum* (Araneae; lower). B: Live specimen of *C. ioanniticus* from Khirbet Haruba cave (Haruva cave). Inset: detail of the median eyes. C: Live specimen of *C. israelensis* from Mimlach cave. Inset: detail of the reduced median eyes. D: Live specimen of *Parasteatoda tepidariorum*, and schematic representation of the expression patterns of paralog pairs of *Ptep-sine oculis* (*soA/soB*), *Ptep-orthodenticle* (*otdA/otdB*), and *Ptep-Optix* (*OptixA/OptixB*) in the eyes. ME: median eyes; ALE: anterior lateral eyes; PLE: posterior lateral eyes; MLE: median lateral eyes; LE: lateral eyes.

The second concern in extending the model derived from pancrustaceans is that a subset of Arachnida exhibits an ancient shared genome duplication, resulting in numerous paralogs of developmental patterning genes. Recent phylogenetic and comparative genomic works on Arachnida have shown that Arachnopulmonata (Ballesteros & Sharma, 2019; Ballesteros, Santibáñez López, Kováč, Gavish-Regev, & Sharma, 2019; Sharma, Kaluziak, Pérez-Porro, González, Hormiga, et al., 2014a), the clade of arachnids that bear book lungs (e.g., spiders, scorpions, whip spiders), retain duplicates of many key transcription factors, such as homeobox genes, often in conserved syntenic blocks (Leite et al., 2018; Schwager et al., 2017; Sharma, Santiago, González-Santillán, Monod, & Wheeler, 2015a; Sharma, Schwager, Extavour, & Wheeler, 2014b). Many of the ensuing paralogs exhibit non-overlapping expression patterns and a small number have been shown to have subdivided the ancestral gene function (subfunctionalization) or acquired new functions (neofunctionalization) (Leite et al., 2018; Paese, Leite, Schönauer, McGregor, & Russell, 2018; Turetzek, Pechmann, Schomburg, Schneider, & Prpic, 2015).

While comparatively little is known about the genetics of arachnid eye development, gene expression surveys of insect retinal determination gene network (RDGN) homologs of two spiders (*Cupiennius salei* and *Parasteatoda tepidariorum*) have shown that this phenomenon extends to the formation of spider eyes as well (Samadi, Schmid, & Eriksson, 2015; Schomburg et al., 2015). Different paralog pairs (orthologs of *Pax6, Six1, Six3, eyes absent, atonal*, *dachshund* and *orthodenticle*) exhibit non-overlapping expression boundaries in the developing eye fields, resulting in different combinations of transcription factor expression in the eye pairs (Samadi et al., 2015; Schomburg et al., 2015). While these expression patterns offer a potentially elegant solution to the differentiation of spider eye pairs, only a few studies with the spider *P. tepidariorum* have attempted to experimentally test the role of these genes in the formation of arachnid eyes. *Ptep-orthodenticle-1* maternal RNA interference (RNAi) knockdown results in a range of anterior defects, including complete loss of the head, which precluded assessment of a role in the formation of the eyes (Pechmann, McGregor, Schwager, Feitosa, & Damen, 2009). *Ptep-dac2* RNAi knockdown results in appendage segment defects, but no eye patterning defects were reported by the authors (Turetzek et al., 2015). More recently, a functional interrogation of *Ptep-Six3* paralogs, focused on labrum development, reported no discernible morphological phenotype, despite a lower hatching rate than controls and disruption of a downstream target with a labral expression domain (Schacht, Schomburg, & Bucher, 2020). Thus, gene expression patterns of duplicated RDGN paralogs have never been linked to eye-related phenotypic outcomes in any arachnopulmonate model. Similarly, the functions of the single-copy orthologs of RDGN genes in groups like mites (Grbić et al., 2007; Telford & Thomas, 1998), ticks (Santos et al., 2013), and harvestmen (Garwood et al., 2014; Sharma, Schwager, Giribet, Jockusch, & Extavour, 2013; Sharma, Tarazona, Lopez, Schwager, Cohn, Wheeler, et al., 2015b) are entirely unexplored, in one case because an otherwise tractable arachnid species lacks eyes altogether (the mite *Archegozetes longisetosus* (Barnett & Thomas, 2012; 2013a; 2013b; Telford & Thomas, 1998).

Investigating the evolution of eye loss in arachnids thus has the potential to elucidate simultaneously (1) the morphogenesis of a poorly understood subset of metazoan eyes (Foelix, 2011; Morehouse et al., 2017), (2) developmental mechanisms underlying a convergent trait (i.e., eye loss in caves) in phylogenetically distant arthropod groups (Protas & Jeffery, 2012; Re et al., 2018), (3) shared programs in eye development common to Arthropoda (through comparisons with pancrustacean datasets) (Cagan, 2009; Stahl et al., 2015; Takagi et al., 2012; ZarinKamar et al., 2011), and (4) the role of ancient gene duplicates in establishing the diversity of eyes in arachnopulmonates (Leite et al., 2018; Samadi et al., 2015; Schomburg et al., 2015).

As first steps toward these goals, we first developed transcriptomic resources for a sister species pair of cave-dwelling *Charinus* whip spiders, wherein one species exhibits typical eye morphology and the other highly reduced eyes (a troglobitic condition). We applied a differential gene expression (DGE) analysis to these datasets to investigate whether candidate RDGN genes with known expression patterns in model spider species (*C. salei*, *P. tepidariorum*) exhibit differential expression in non-spider arachnopulmonates, as a function of both eye condition and developmental stage. To link bioinformatic inference of expression patterns with functional outcomes, we interrogated the function of three candidate RDGN genes identified from DGE in a model arachnopulmonate, using RNAi in the spider *P. tepidariorum*, which exhibits the same number and types of eyes as whip spiders. We provide functional evidence that one of these candidates, *sine oculis/Six1*, is necessary for the development of all spider eye types.

## Results

### *Charinus ioanniticus* and *Charinus israelensis* embryonic transcriptomes

As an empirical case of closely related, non-spider arachnopulmonate sister species pair that constitutes one epigean and one troglobitic species, we selected the whip spider species *Charinus ioanniticus* and *C. israelensis* (Fig. 1 B–C). Whip spiders, arachnopulmonates of the order Amblypygi, are commonly found in cave habitats ranging from rain forests, savannahs and deserts (Weygoldt, 2000). The recently described troglobitic species *Charinus israelensis* (reduced-eyes) occurs in close proximity to its congener *Charinus ioanniticus* (normal-eyes) in caves in the Galilee, northern Israel (Miranda et al. 2016). Given that the formation of Levantine cave refuges is considerably recent, *C. israelensis* and *C. ioanniticus* are likely sister species with a small time of divergence, an inference supported by their similar morphology (Miranda et al 2016). We collected ovigerous females from both species in caves in Israel and extracted RNA from embryos (SI Appendix, Table S1). Embryos of whip-spiders (*Phrynus marginemaculatus*) achieve a deutembryo stage around 20–25 days after egg laying (dAEL), a stage where most external features of the embryo, such as tagmosis and appendages are fully formed, but not the eyes (Weygoldt, 1975). The deutembryo hatches from the egg membrane inside the broodsac carried by the mother, but remains in this stage relatively unchanged for around 70 days. The eyes begin to form around 50 dAEL, but the eye spots become externally visible and pigmented only close to hatching (90 dAEL) (Weygoldt, 1975).

For *de novo* assembly of the embryonic transcriptomes of *C. ioanniticus* and *C. israelensis*, we extracted RNA from all embryonic deutembryo stages collected in the field (see Supplementary Information; table 1 for localities and sample explanations). Assemblies include two deutembryo stages before eyespot formation and one deutembryo stage bearing eyespots for *C. ioanniticus*; and two early deutembryo stages for *C. israelensis* (SI Appendix, Fig. S1).

The assembly of *C. ioanniticus* reads resulted in 219,797 transcripts composed of 143,282,365 bp with and N50 of 1122 bp (more than 50% of transcripts are 1122 bp or longer) (SI Appendix, Table S2). Universal single copy ortholog benchmarking with BUSCO v3.0 (Waterhouse et al., 2017) indicated 93.8% completeness, with 5.7% of BUSCO genes exhibiting duplication.

The assembly *C. israelensis* resulted in a higher number of transcripts: 663,281 transcripts composed of 230,044,656 bp and with N50 of 1045 bp. The BUSCO analysis shows 95.2% completeness, which is similar to the value for *C. ioanniticus* assembly.

### RDGN gene duplication in *Charinus* whip spiders

Amblypygi is inferred to be nested stably in Arachnopulmonata, the clade of arachnids that bear book lungs (Ballesteros & Sharma, 2019; Giribet, 2018; Lozano-Fernandez et al., 2019; Rota-Stabelli et al., 2010; Sharma, Kaluziak, Pérez-Porro, González, Hormiga, et al., 2014a). Recent evidence suggests that the common ancestor of arachnopulmonates has undergone a whole- or partial-genome duplication affecting large gene families, such as homeobox genes (Leite et al., 2018; Schwager et al., 2017; Sharma, Santiago, González-Santillán, Monod, & Wheeler, 2015a). The stable phylogenetic position of Amblypygi in Arachnopulmonata predicts that genes in RDGN that are duplicated in spiders, should also be duplicated in *Charinus* whip spiders. To test this hypothesis, we performed phylogenetically-informed orthology searches on the newly assembled embryonic transcriptomes of both *Charinus* species, and conducted phylogenetic analysis with orthologs across selected arthropod species. We discovered that homologs of *atonal, Pax6, dachshund, sine oculis* (*Six1*), *Optix* (*Six3*), and *orthodenticle* are duplicated in *Charinus*, whereas *eyegone* and *eyes absent* occur as single-copy orthologs (these latter two also occurring single-copy in spiders) (Fig. 2).

**Figure 2:**
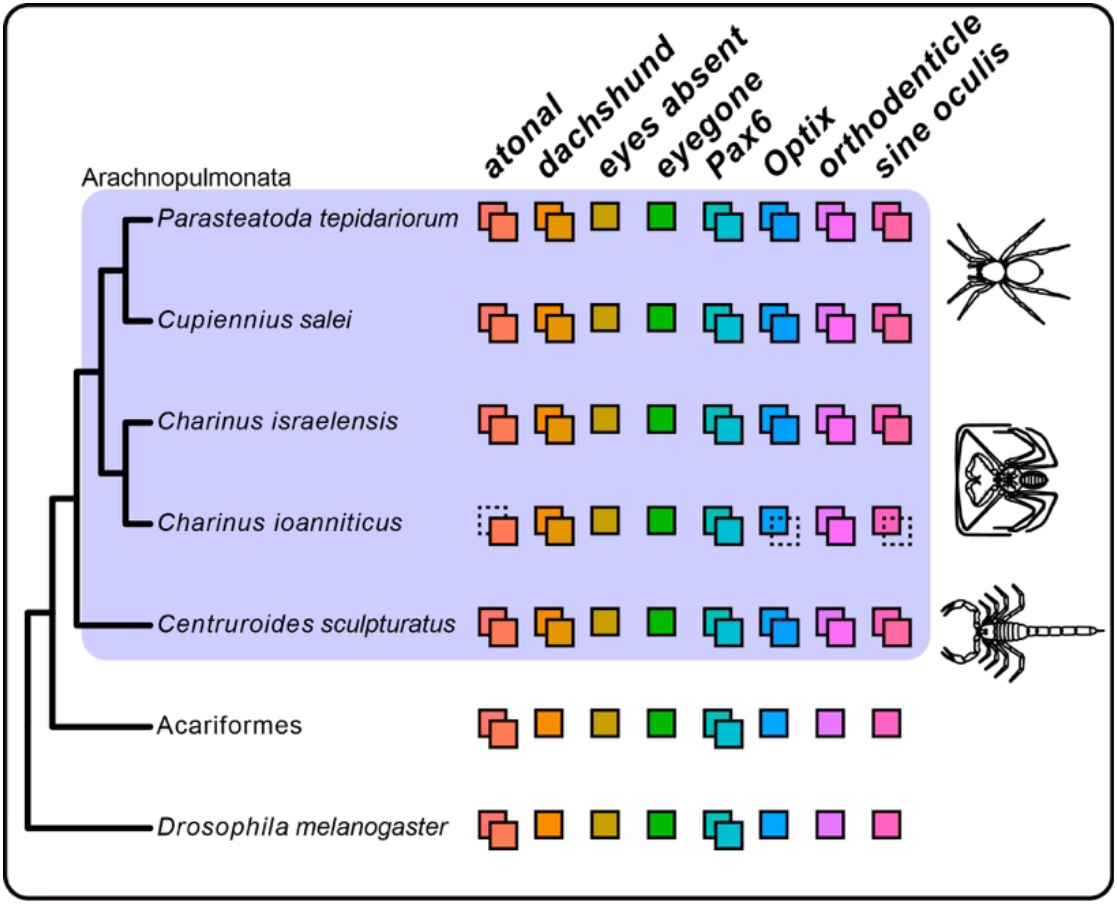
Phylogenetic distribution of Retinal Determination Gene Network (RDGN) genes in an insect (*Drosophila melanogaster*), a non-arachnopulmonate arachnid group (Acariformes: *Dinothrombium tinctorium*; *Tetranychus urticae*) and Arachnopulmonata (spider: *Parasteatoda tepidariorum*; scorpion: *Centruroides sculpturatus*), including newly discovered orthologs in *Charinus* whip spiders (Amblypygi). Colored squares indicate number of gene copies for each RDGN gene. Dotted squares indicate missing data, not gene loss. For comprehensive list of duplicated genes in Arachnopulmonata see Schwager et al. (2017) and Leite et al. 2018. Gene trees and alignments for each gene are available in SI Appendix Dataset S1.

#### atonal

The *atonal* gene tree showed poor resolution (SI Appendix, Fig. S2), hampering unambiguous assignment of the whip spider genes to *atonal* copies previously annotated in spiders (Samadi et al., 2015; Schwager et al., 2017). *D. melanogaster* copies of *atonal* and *amos* clustered together forming a clade with other pancrustacean and myriapod sequences, suggesting these paralogs are restricted to Mandibulata. The fruit fly *cousin of atonal* (*cato*) formed a clade including the *Cupiennius salei* sequence of *atonalB* whereas the second copy of *C. salei*, *atonalA*, is found in an independent clade with only arachnid sequences. It is in this later clade that the only sequences of *Charinus* related to *atonal* are found, in turn forming two separate clades with clear amino acid differences between these copies (SI Appendix, Dataset S1; *atonal* alignment). Herein, these copies are labeled *atonalA* (*atoA*) and *atonalB* (*atoB*). Note that the reference genomic sequences, annotated as “*atonal like homolog 8 like*” (Ptep XP 0159181091), is found orthologous to the gene *net* in *D. melanogaster*.

#### Pax6

In *D. melanogaster*, there are two paralogous copies of the vertebrate *Pax6*, *eyeless* and *twin of eyeless*. This duplication seems to be shared across all arthropods and both *Pax6* copies have been characterized in spiders (Samadi et al., 2015; Schomburg et al., 2015). The gene tree of *Pax6* homologues clearly identified a clade for *toy* including chelicerate and mandibulate copies, but no *Charinus* sequences are found in this clade (SI Appendix, Fig. S3). The sister clade (*eyeless*) consists only of pancrustacean sequences whereas the chelicerate copies, previously annotated as *eyeless* orthologs, are found in a separate clade. Among these, two distinct genes, herein dubbed *Pax6A* and *Pax6B*, are present in both *Charinus* species. Sequence similarity searches (blastp) of both *Pax6A* and *Pax6B* against the genome of *Drosophila melanogaster* points to *Dmel-toy* as the best hit, followed by *Dmel-ey*. Therefore, although the homology of these copies with *Dmel-ey/toy* is evident, it is not trivial to assign these to either of these genes or if these represent taxon-restricted duplicates of *eyeless*.

#### eyegone/twin of eyegone

These members of the Pax gene family are paralogous in *D. melanogaster* but occur as single copy in arachnids. Single copy orthologs of eyg/toe are present in the two target Amblypygi species (SI Appendix, Fig. S3).

#### dachshund

Spiders and scorpions have two paralogous copies of *dachshund* (Nolan, Santibáñez López, & Sharma, 2020; Turetzek et al., 2015). Two copies are present in the transcriptomes of both *Charinus* species and are here termed *dacA* and *dacB* (SI Appendix, Fig. S4). The *C. israelensis dacB* is assembled in two different gene fragments that overlap by three amino acids (SI Appendix, Fig. S4; see *dachshund* alignment in SI Appendix Dataset S1). The *C. ioanniticus dacA* copy is also assembled as two different gene fragments with little sequence overlap but being part of the *dacA* clade (SI Appendix, Fig. S4).

#### eyes absent

This single-copy orthologs are found in arthropods and arachnids alike and is represented in both *Charinus* species. The association of transcript to this gene is unambiguous for both amblypigid species (SI Appendix, Fig. S5).

#### orthodenticle

As with spiders, there are two copies homologous to *Dmel-otd* in *Charinus.* The resolution of the gene tree is poor and does not allow uncontroversial association to spider orthologs (SI Appendix, Fig. S6). *Charinus* copies are termed *otdA* and *otdB*.

#### Optix

There are two very similar copies of *Optix* in *C. israelensis* and one in *C. ioanniticus* (SI Appendix, Fig. S7). The *C. ioanniticus* copy is termed *OptixA*. The two copies of *C. israelensis* show very conserved amino acid sequences but clear nucleotide differences. Although the gene tree with the reference genome shows them more closely allied to one of the spider paralogous copies of *Optix* (Ptep NP 00130752.1), a reduced analysis including *Cupiennius salei* and *P. tepidariorum* copies, suggests that the *Charinus* copies are independent duplications. Here the two whip spider copies are dubbed *OptixA* and *OptixB* but they should not be considered orthologous to the spider *OptixA/B*.

#### sine oculis

Two copies of *sine oculis* are found in *C. israelensis* and one in *C. ioanniticus*. Both copies are nested in a clade with *Ptep-soA* (SI Appendix, Fig. S8). These are herein dubbed as *soA* and *soB* given that orthology with either spider copy is unclear.

### RDGN genes in whip spider eye formation: comparing early and late stages of *C. ioanniticus*

The expression of paralog pairs of *Pax6*, *sine oculis*, *Optix*, *eyes absent*, *atonal*, *dachshund* and *orthodenticle* in the developing eyes of the spiders (Samadi et al., 2015; Schomburg et al., 2015), and the occurrence of the same paralogs in *Charinus* whip spiders, suggest that these genes may also be involved in the formation of eyes in whip spiders. We investigated this idea by comparing the expression levels of these RDGN genes in the stages before eye-spot formation versus a stage after eye-spot formation in the eye-bearing whip spider *Charinus ioanniticus* (henceforth “Comparison 1”; Fig. 3A).

**Figure 3:**
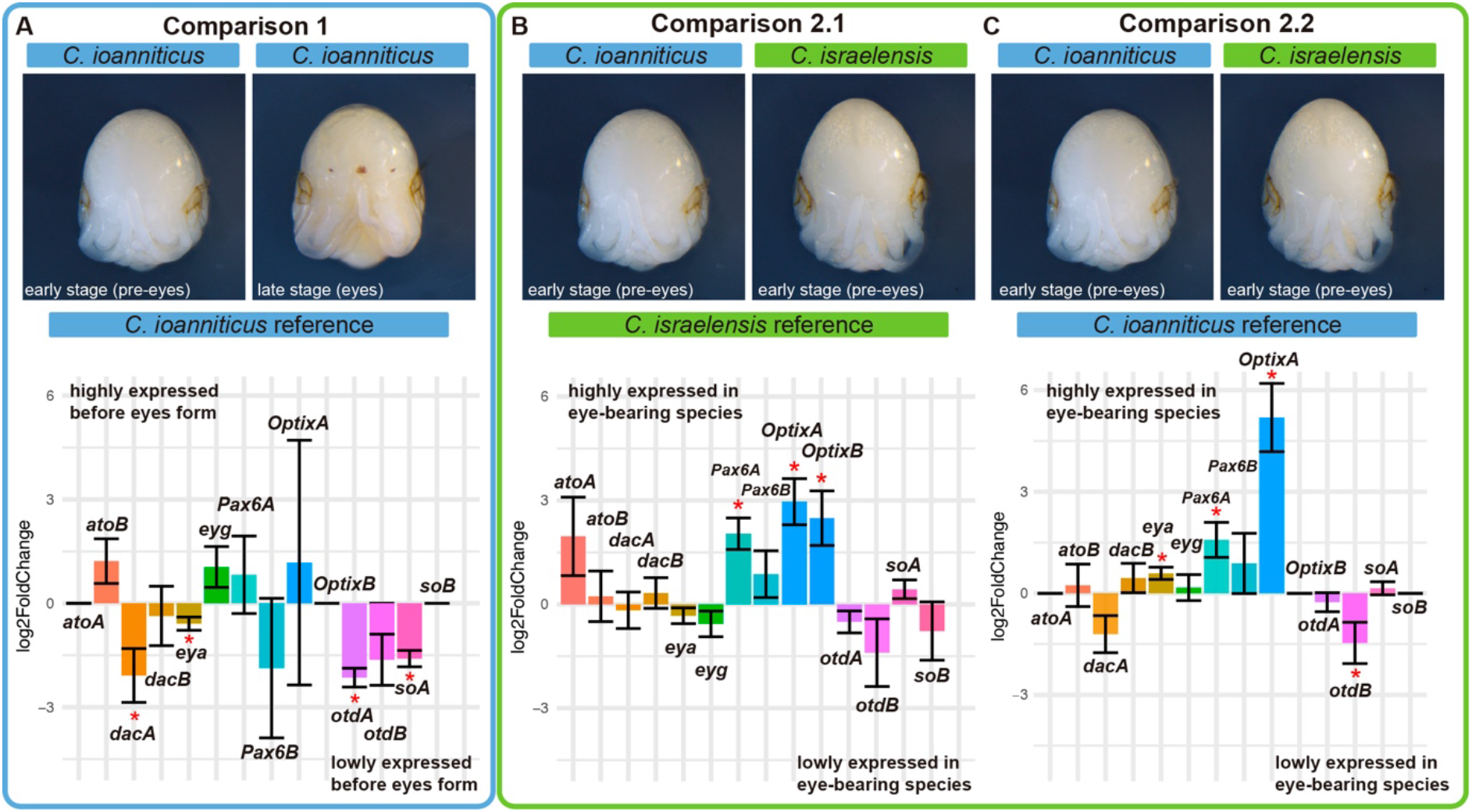
Differential gene expression analysis of Retinal Determination Gene Network (RDGN) genes in *Charinus* whip spider deutembryos. Bar graphs display log2 fold change of selected RDGN genes. Direction of differential gene expression always follows sample to the left. A: Comparison 1; Comparison between reads of early (pre-eyespot) and late deutembryos (eyespot) of the eye-bearing species *C. ioanniticus* mapped onto *C. ioanniticus* transcriptome. B: Comparison 2.1; Comparison between reads of early deutembryo of *C. ioanniticus* and early deutembryo of *C. israelensis* mapped onto *C. israelensis* transcriptome. C: Comparison 2.2; Comparison between reads of early deutembryo of *C. ioanniticus* and early deutembryo of *C. israelensis* mapped onto *C. ioanniticus* transcriptome. *atoA/B*: *atonalA/atonalB*; *dacA/B*: *dachshundA/B; eya: eyes absent; eyg: eyegone; otdA/B: orthodenticleA/B; soA/B: sine oculisA/B*. Asterisks denote genes that were differentially expressed with a p_adj_>0.05. Log_2_FC = 0 for *atoA, OptixB*, and *soB* for Comparison 1 and Comparison 2.2 are due to the absence of those paralogs in *C. ioanniticus* reference transcriptome.

We mapped reads of both treatments to the reference transcriptome of *C. ioanniticus* using the quasi-alignment software Salmon v. 1.1.0 (Patro, Duggal, Love, Irizarry, & Kingsford, 2017) and conducted a differential gene expression analysis of Comparison 1 using DESeq2 v 1.24.0 (Love, Huber, & Anders, 2014) (SI Appendix, Fig. S9). These comparisons showed that *Cioa-dacA*, *Cioa-otdA*, *Ciao-eya* and *Cioa-soA* are significantly over-expressed (p_adj_ < 0.05) in the eyespot stage in comparison with the stage before eyespot formation (Fig. 3A). While we cannot rule out that the differences in gene expression are due to other developmental differences between the two stages sequenced, these results highlighted these four RDGN genes as promising candidates involved in the formation of eyes in whip spiders.

### RDGN genes in whip spider eye reduction: comparing *C. ioanniticus* and *C. israelensis*

Blindness in adults of the model cave fish *Astyanax mexicanus* is a result of an embryonic process in which the rudimentary eye of the embryo is induced to degenerate by signals emitted from the lens tissue (Jeffery, 2009). Both early and late expression of RDGN genes, such as Pax6, are responsible for the reduction of eyes in fish from cave populations (Jeffery, 2009; Strickler, Yamamoto, & Jeffery, 2001). Likewise, in the isopod crustacean *Asellus aquaticus* cave blindness has a strong genetic component and mechanisms of eye reduction also act at embryonic stages (Mojaddidi, Fernandez, Erickson, & Protas, 2018; Protas et al., 2011). The embryonic development of the reduced-eyes whip spider *C. israelensis* has not been explored to date, but we expect that reduction of eyes results from changes in embryonic gene expression during the deutembryo stage (Weygoldt, 1975). We investigated this possibility by quantifying the relative gene expression of RDGN genes in comparable embryonic stages of *C. israelensis* (reduced eyes) and *C. ioanniticus* (normal eyes) embryos before eye-spot formation (SI Appendix Table S1; Figure S1). Using the DGE approach from Comparison 1, we conducted a heterospecific analysis using as the reference either the *C. israelensis* transcriptome (henceforth “Comparison 2.1”) or the *C. ioanniticus* transcriptome (henceforth “Comparison 2.2”).

Both analyses are anchored on the premise that a hybrid mapping between the sister species is possible given the recent divergence between them. The mapping rate of the *C. ioanniticus* reads was similar regardless of the reference species, (96.74% and 96.59% respectively for *C. ioanniticus* and *C. israelensis*). In the case of the reads from *C. israelensis* embryos, mapping rate to the conspecific (96.8%) transcriptome was higher than when mapping against *C. ioanniticus* (82.45%). The similar mapping rate of *C. ioanniticus* reads suggests that the two whip spiders are sufficiently closely related to generate interspecific comparisons of gene expression. Comparisons 2.1 and 2.2 yielded similar results with respect to the direction of differentially expressed RDGN genes (Fig. 3B–C). In comparison 2.1, *Pax6A*, *OptixA* and *OptixB* are significantly over-expressed in the normal-eyes species, with expression levels at least 4 times higher than in the reduced-eyes species (log_2_FC > 2; p_adj_ < 0.05) (Fig. 3B; SI Appendix, Fig. S10). In comparison 2.2, *Pax6A* and *OptixA* are also over-expressed in *C. ioanniticus* (p_adj_ < 0.05), and so is *eyes absent* (p_adj_ < 0.05; Fig. 3C). In comparison 2.2, *orthodenticle-B* appears under-expressed in the normal-eyes species (p_adj_ < 0.05 (Fig. 3C; SI Appendix, Fig. S11). We note that the magnitude of log_2_FC and significance values differed considerably between analysis. Nonetheless, *Pax6A* and *OptixA* were consistently over expressed in the normal-eyes species, highlighting these two genes as promising candidates involved in the reduction of eyes in *Charinus israelesis*.

### *sine oculis* is necessary for principal and secondary eye development in a model arachnopulmonate

Our bioinformatic analysis in the whip spider system suggested that *eyes absent* and paralogs of *sine oculis*, *orthodenticle*, and *dachshund* may be involved in the normal formation of eyes in *C. ioanniticus* (Comparison 1). We also found evidence that *Pax6* and a paralog of *Optix* may be involved in the reduction of eyes in the cave whip spider *C. israelensis*. To link bioinformatic reconstructions of gene expression with functional outcomes, we interrogated the function of RDGN genes using parental RNA interference (RNAi) in the spider *Parasteatoda tepidariorum*. We selected *Ptep-soA* (*Ptep-so1 sensu* Schomburg et al. 2015), *Ptep-otdB* (*Ptep-otd2 sensu* Schomburg et al. 2015) and *Ptep-OptixB (Ptep Six3.2 sensu* Schomburg et al. 2015). In *P. tepidariorum*, these genes are known to be expressed in all eye types, in the median eyes only, and in the lateral eyes, respectively (Fig. 1D) (Schomburg et al., 2015).

Early expression of *Ptep-soA* is detected in lateral domains of the head lobes (stage 10) corresponding to the principal and secondary eyes, and continues until the pre-hatching stage 14 (Schomburg et al., 2015). Expression of *Ptep-soA* on wild type stage 14.1 embryos is bilaterally symmetrical on all eyes and uniformly strong (Fig. 4A–B). By stage 14.2, it remains strong on the principal eyes but it is stronger at the periphery of the secondary eye spots (Fig. 4A, C).

**Figure 4:**
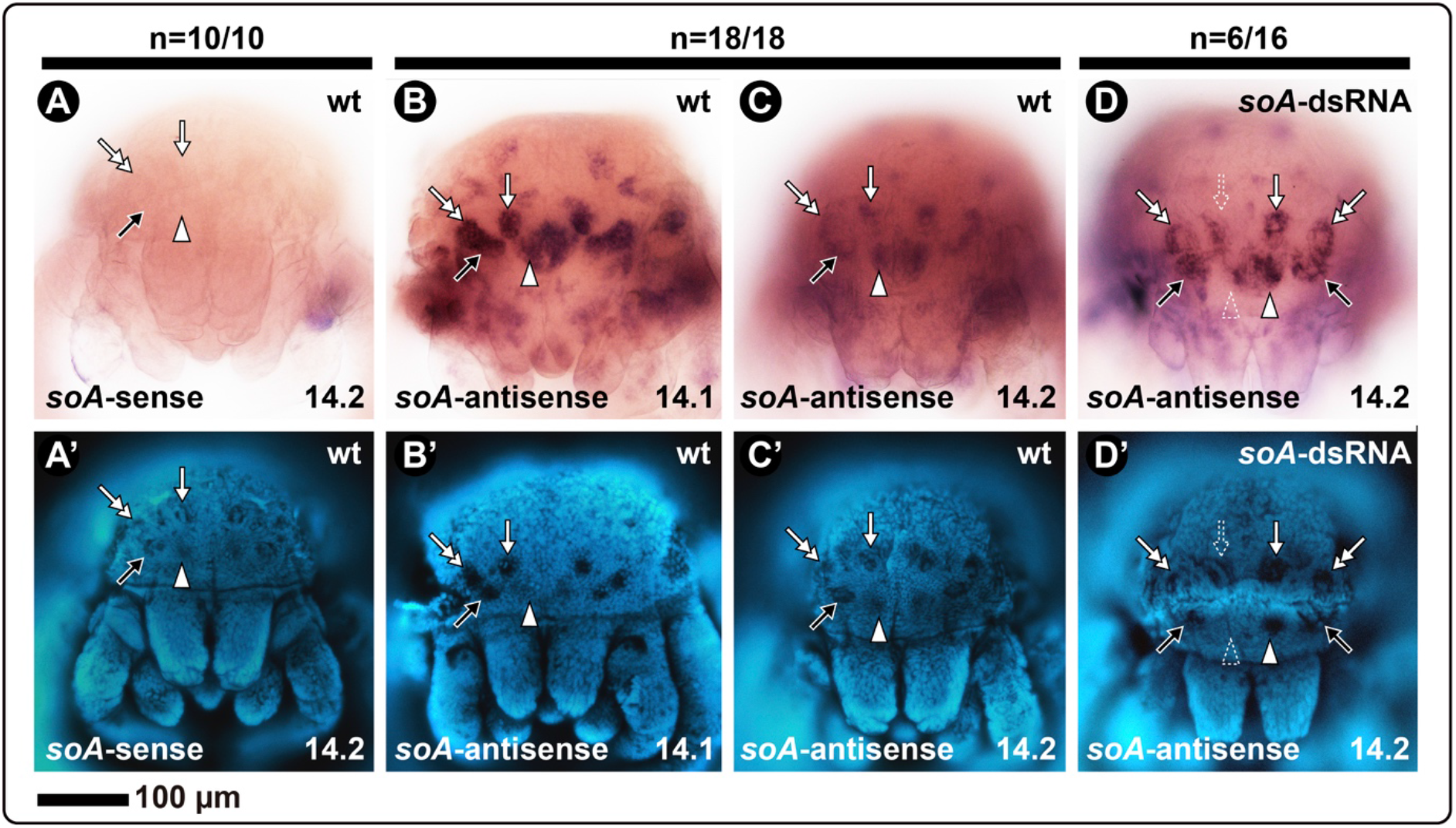
In situ hybridization using DIG-labeled riboprobes for *Ptep-soA* in late embryos of the spider *Parasteatoda tepidariorum*. All embryos in frontal view. A–D: bright field images. A’–D’: Same embryos, in Hoechst staining. A: Sense probe of a stage 14.2 embryo (no signal). B: Antisense probe on a wild type stage 14.1 embryo. C: Antisense probe on a wild type stage 14.2 embryo. D: Antisense probe on a stage 14.2 embryo from the *Ptep-soA* dsRNA-injected treatment. *soA*: *sine oculis A*. White arrowhead: median eye; Black arrow: anterior lateral eye; White arrow: median lateral eye; Double white arrow: Posterior lateral eye. Dotted arrowhead/arrow indicate asymmetrical expression and eye defect. Sample sizes are indicated above each treatment.

*P. tepidariorum* hatchlings, or postembryos, initially have no externally visible lenses and pigment. The red pigment and lenses of all eyes, and the reflective tapetum of the lateral eyes, become progressively recognizable in the 48 hours (at 26°C) until the animal molts into the first instar with fully formed eyes (SI Appendix, Video S1) (see also Mittmann & Wolff, 2012). We fixed embryos from *Ptep-soA* dsRNA-injected and dH2O-injected treatments between 24h-48h, which encompasses stages where the eyes of postembryos are already recognizable until the first instar.

Negative control experiments (dH2O-injected females) yielded postembryos with eye morphology indistinguishable from wild type animals: the median eyes (ME; principal eyes) have an inferior semi-lunar ring of red pigment and lack the tapetum; and all pairs of lateral eyes (secondary eyes) have the canoe-shaped tapetum type (Foelix, 2011; Land, 1985), which is split in the middle and surrounded by red pigment (Fig. 5A; panel 1). We observed misshaped tapeta on the lateral eyes of some postembryos on the earlier side of the developmental spectrum of fixed animals, but that was never observed on postembryos close to molting or first instars (SI Appendix, Fig. S12). It is unclear if this reflects a natural variation of early developing tapetum or an artifact of sample preparation.

**Figure 5:**
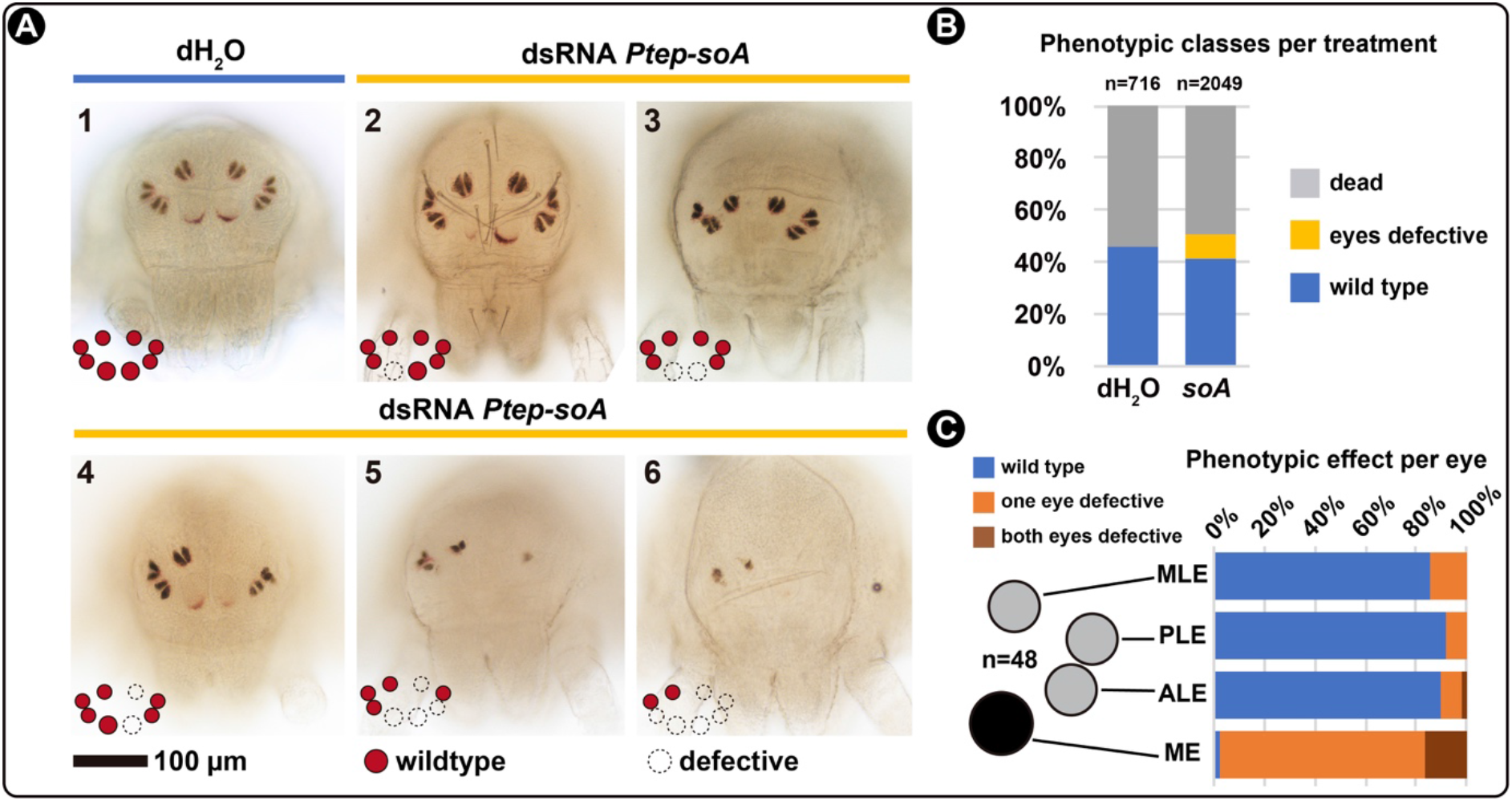
RNA interference against *Ptep-sine oculis A*. A: Bright field images of the spider *Parasteatoda tepidariorum* postembryos resulting from control treatment (dH_2_O-injected, panel 1) and double stranded RNA (dsRNA) injected treatment (panels 2–6), in frontal view. B: Frequencies of each phenotypic class per treatment from the combined clutches of all females. See SI Appendix Fig. S13 for counts per clutch. C: Frequencies of symmetrical, asymmetrical, and wild type eyes quantified from a subset of 48 individuals with eye reduction phenotype. See SI Appendix Fig. S12 for figures of all specimens and coding, and Material and Methods for the scoring criteria. ME: median eyes; ALE: anterior lateral eyes; PLE: posterior lateral eyes; MLE: median lateral eyes. Schematics for the different eye types follows the nomenclature in Figure 1.

Embryos from *Ptep-soA* dsRNA-injected females are also able to hatch into postembryos and continue molting to adulthood (SI Appendix, Video S2). However, a subset of the embryos of dsRNA-injected treatment (9.5%; n=195/2049) exhibits a spectrum of eye defects that was not observed on the controls (Fig. 5A–B; SI Appendix, Fig. S13). The defects occurred on all eyes, namely medium eyes (ME), anterior lateral eyes (ALE), posterior lateral eyes (PLE), and medium lateral eyes (MLE) (Fig. 5A). Affected medium eyes have reduced pigmentation or complete absence (Fig. 5A, panels 2–6), while lateral eyes also exhibited defects of the tapetum or complete absence of the eye (Fig. 5A, panels 4–6).

We selected a subset of the knockdown postembryos initially scored as having any eye defect (n=48) for quantifying the degree of effect per eye type, and the proportion of symmetrical and mosaic eye phenotypes in our sample. Medium eyes are affected in almost all cases (97%), whereas the three lateral eye types were similarly lowly affected (MLE: 14%; PLE: 8%; ALE: 10%) (Fig. 5C; SI Appendix, Fig. S12; detailed scoring criteria in Material and Methods). The majority of defective eyes are mosaics, meaning that a given eye pair is affected only on one side of the animal (Fig. 5C; SI Appendix, Fig. S12).

Parental RNAi against *Ptep-soA* did not completely abolish its expression, as detected by in situ hybridization (Fig. 4D; see Material and Methods). Nevertheless, we detected asymmetrical reduction of *Ptep-soA* expression on single eyes of a subset of stage 14 embryos (n=6/16; Fig. 4D), which closely correlates with the predominance of mosaic phenotypes observed in late postembryos (Fig. 5C).

Parental RNAi experiments using the same protocol targeting *Ptep-otdB* and *Ptep-OptixB* did not result in any detectable phenotypic effects on the eyes of embryos from dsRNA-injected treatment (two and six females injected, respectively; counts not shown). These results accord with a recent study that knocked down both *Optix* paralogs *P. tepidariorum* and did not recover eye defects (Schacht et al., 2020)

## Discussion

### Paralogy of RDGN members in arachnopulmonates

Amblypygi have a critical placement within arachnid phylogeny, as they are part of a trio of arachnid orders (collectively, the Pedipalpi, comprised of Amblypygi, Thelyphonida, and Schizomida), which in turn is the sister group to spiders. Whereas the eyes of spiders have greatly diversified in structure, function, and degree of visual acuity (particularly the eyes of hunting and jumping spiders), the arrangement and number of eyes in Amblypygi likely reflects the ancestral condition across Tetrapulmonata (= spiders + Pedipalpi), consisting of three pairs of simple lateral ocelli and a pair of median ocelli; a similar condition is observed in basally branching spider groups like Mesothelae and Mygalomorphae, as well as Thelyphonida (vinegaroons). However, while developmental genetic datasets and diverse genomic resources are available for spiders and scorpions (Oda & Akiyama-Oda, 2020; Posnien et al., 2014; Schwager et al., 2017; Sharma, Schwager, Extavour, & Wheeler, 2014b), the developmental biology of the other three arachnopulmonate orders has been virtually unexplored in the past four decades beyond a single work describing the embryology of one North American amblypygid species (Weygoldt, 1975). To address this gap, we focused our investigation on a sister species pair of cave whip spiders and generated the first embryonic transcriptomes for this order. These datasets are immediately amenable to testing the incidence of RDGN duplicates previously known only from two spiders (Samadi et al., 2015; Schomburg et al., 2015) and their putative effects in patterning eyes across Arachnopulmonata broadly.

The inference of a partial or whole genome duplication (WGD) in the most recent common ancestor of Arachnopulmonata is supported by the systemic duplications of transcription factors and synteny detected in the genomes of the scorpion *Centruroides sculpturatus*, and the spider *Parasteatoda tepidariorum*, as well as homeobox gene duplications detected in the genome of the scorpion *Mesobuthus martensii* and transcriptome of the spider *Pholcus phalangioides* (Leite et al., 2018; Schwager et al., 2017). Additional evidence comes from shared expression patterns of leg gap gene paralogs in a spider and a scorpion (Nolan et al., 2020). Embryonic transcriptomes are particularly helpful in the absence of genomes, as several duplicated genes, such as some homeobox genes, are only expressed during early stages of development (Leite et al., 2018; Sharma, Santiago, González-Santillán, Monod, & Wheeler, 2015a; Sharma, Schwager, Extavour, & Wheeler, 2014b). Our analysis of *Charinus* embryonic transcriptomes shows that RDGN gene duplicates observed in spiders also occur in whip spiders, supporting the hypothesis that these paralogous copies originated from a shared WGD event in the common ancestor of Arachnopulmonata.

The conservation of some transcription factors patterning eyes is widespread in the Metazoan tree of life (Vopalensky & Kozmik, 2009). In the model fruit fly *D. melanogaster*, the homeobox Pax6 homolog, *eyeless*, was the first transcription factor identified as a “master gene”, necessary for compound eye formation and capable of inducing ectopic eye formation (Gehring & Ikeo, 1999; Kumar, 2009). The Pax6 protein is essential for eye formation across several metazoan taxa, which has fomented ample debate about the deep homology of gene regulatory networks in patterning structurally disparate eyes (Carroll, 2008; Shubin, Tabin, & Carroll, 2009; Vopalensky & Kozmik, 2009). In the case of *sine oculis* (Six1/2), orthologs are found across metazoans (Bebenek, Gates, Morris, Hartenstein, & Jacobs, 2004; Byrne et al., 2017; Rivera et al., 2013). Evidence that *sine oculis* is required for the eye patterning in other bilaterians comes from expression pattern in the developing eyes of the annelid *Platynereis dumerilii* (Arendt, Tessmar, Medeiros de Campos-Baptista, Dorresteijn, & Wittbrodt, 2002), and functional experiments in the planarian *Girardia tigrina* (Pineda et al., 2000). Therefore, studies interrogating the genetic bases of eye formation in chelicerate models have the potential to clarify which components of the eye gene regulatory network of Arthropoda evolved in the MRCA of the phylum, and which reflect deep homologies with other metazoan genes.

### A conserved role for a *sine oculis* homolog in patterning arachnopulmonate eyes

The eyes of arthropods are diverse in number, arrangement, structure and function (Paulus, 1979). Both types of eyes observed in Arthropoda, the faceted eyes (compound) and single-lens eyes (ocelli), achieve complexity and visual acuity in various ways. To mention two extremes, in Mandibulata the compound eyes of mantis shrimps (Stomatopoda) achieve a unique type of color vision and movements by using 12 different photoreceptive types and flexible eye-stalks (Daly, How, Partridge, & Roberts, 2018; Marshall, Cronin, & Kleinlogel, 2007; Thoen, How, Chiou, & Marshall, 2014). In Arachnida, the simple-lens median eyes of some jumping spiders (Salticidae) have exceptional visual acuity in relation to their eye size, achieve trichromatic vision through spectral filtering, and can move their retina using specialized muscles (Harland, Li, & Jackson, 2012; Land, 1985; Zurek et al., 2015). Comparative anatomy suggests that the common ancestor of Arthropoda had both lateral compound eyes and median ocelli that then became independently modified in the arthropod subphyla (Morehouse et al., 2017; Paulus, 1979). While in situ hybridization data for selected RDGN genes across arthropods generally support the hypotheses of eye homology, comparative developmental datasets remain phylogenetically sparse outside of Pancrustacea (Samadi et al., 2015; Schomburg et al., 2015)

We therefore applied a bioinformatic approach in a study system that lacked any genomic resources (Amblypygi) to assess whether RDGN homologs are transcriptionally active during the formation of eyes in the eye-bearing *C. ioanniticus* (Comparison 1), as well as those that may be putatively involved in eye loss in its troglobitic sister species (Comparison 2). As first steps toward understanding how arachnid eyes are patterned, our experiments demonstrated that *soA*, a *sine oculis* paralog identified as differentially expressed during the formation of eyes in *C. ioanniticus*, is necessary for patterning all eyes of a model arachnid system with the same eye configuration (*Parasteatoda tepidariorum*). Thus, we provide the first functional evidence that part of the RDGN is evolutionarily conserved in the most recent common ancestor (MRCA) of insects and arachnids, and by extension, across Arthropoda.

The advantage of such a bioinformatic approach is that it can potentially narrow the range of candidate genes for functional screens, due to the inherent challenges imposed by paralogy when assessing gene function. Eye reduction in the cave fish *Astyanax mexicanus* has been shown to involve differential expression of genes known to be involved in eye patterning in model organisms, such as *hedgehog* and *Pax6* (*eyeless/toy*) (Jeffery, 2009; Protas & Jeffery, 2012). In addition, other “non-traditional” candidates have been identified, such as *hsp90* (Jeffery, 2009). Likewise, evidence from quantitative trait loci mapping in cave populations of the troglobitic crustacean *Asellus aquaticus* shows that eye loss phenotype is correlated with loci that are not part of the RDGN (Protas et al., 2011; Protas & Jeffery, 2012). The results of the DGE analysis in whip spiders underscore the potential of a DGE approach to triangulate targets among candidate genes in non-model species more broadly. Future efforts in the *Charinus* system should focus on dissecting individual eye and limb primordia of embryos of both species, in order to identify candidate genes putatively involved in the reduction of each eye type, as well as compensatory elongation of the sensory legs of the troglobitic species, toward downstream functional investigation.

### Do gene duplications play a role in the functional diversification of arachnopulmonate eyes?

A challenge in studying arachnopulmonate models to understand ancestral modes of eye patterning in Arthropoda is the occurrence of RDGN duplicates in this lineage. Our orthology searches and phylogenetic analysis showed that the evolutionary history of genes is not always resolved using standard phylogenetic methods, as short alignable regions and/or uncertainty of multiple sequence alignments can result in ambiguous gene trees. One way to circumvent this limitation is by analyzing expression patterns via in situ hybridization between paralogs in different arachnids in order to determine which patterns are plesiomorphic (Leite et al., 2018; Nolan et al., 2020; Turetzek et al., 2015). Nonetheless, the possibility of subfunctionalization and neofunctionalization may also complicate such inferences because discerning one process from the other is analytically challenging (Sandve, Rohlfs, & Hvidsten, 2018).

Genetic compensation of gene paralogs is another confounding variable, which can be accounted for by experimental advances in model organisms (e.g., Shull et al., 2020). We note that the overall penetrance in this experiment is low (9.5%) when compared to some studies in *P. tepidariorum* (e.g., Khadjeh et al 2012; >59% in *Ptep-Antp* RNAi). Wide variance in penetrance has been reported by several research groups in this system, with phenotypic effects varying broadly even within individual experiments (e.g., Fig. 5 of Akiyama-Oda & Oda, 2006; Fig. S5 of Schwager, Pechmann, Feitosa, McGregor, & Damen, 2009). Furthermore, some genes have empirically proven intractable to misexpression by RNAi in *P. tepidariorum*, with one case suggesting functional redundancy of posterior Hox genes to be the cause (Khadjeh et al., 2012). Double knockdown experiments have been shown to exhibit poor penetrance (0-1.5%) in *P. tepidariorum* as well (Fig. S3 of Khadjeh et al. 2012; Fig. S1 of Setton et al., 2017), and to our knowledge, no triple knockdown has ever been achieved. While we cannot rule out functional redundancy with other RDGN paralogs in the present study, the low penetrance we observed may also be partly attributable to our conservative phenotyping strategy (see Material and Methods), which did not assess a possible delay in eye formation and emphasized dramatic defects in eye morphology for scoring.

The occurrence of RDGN gene duplications in Arachnopulmonata, in tandem with improving functional genetic toolkits in *P. tepidariorum* (e.g., Pechmann, 2016), offers a unique opportunity of studying the role of sub- and neofunctionalization in the development of their eyes, and a possible role of this process in the diversification of number, position and structure of the eyes in an ancient group of arthropods (Harland et al., 2012; Land, 1985; Morehouse et al., 2017; Paulus, 1979; Zurek et al., 2015). The genomes of mites, ticks, and harvestmen (Grbić et al., 2011; Hoy et al., 2016) (Gulia-Nuss et al., 2016) reveal that apulmonate arachnid orders have not undergone genome duplication events as seen in Arachnopulmonata (Schwager et al., 2017), or horseshoe crabs (Kenny et al., 2015; Nossa et al., 2014; Zhou et al., 2020). Future comparative studies focused on understanding the ancestral role of chelicerate RDGN genes should additionally prioritize single-copy orthologs in emerging model systems independent of the arachnopulmonate gene expansion, such as the harvestman *Phalangium opilio* (Sharma et al., 2013; Sharma, Schwager, Extavour, & Giribet, 2012).

## Materials and Methods

### Animal collection

Three ovigerous females of the normal-eyes species, *Charinus ioanniticus* (ISR021-2; ISR021-3; ISR021-4), and two egg-carrying females of the reduced-eyes species, *Charinus israelensis* (ISR051-4; ISR051-6), were hand collected in caves in Israel in August 2018 (Supplementary Information; Table 1). Females were sacrificed and the brood sacs containing the embryos were dissected under phosphate saline buffer (PBS). For each female, a subset of the embryos (5 to 13 individuals) was fixed in RNAlater solution after poking a whole into the egg membrane with fine forceps, while the remaining embryos of the clutch were fixed in a 4% formaldehyde/PBS solution to serve as vouchers (SI Appendix, Table S1). Adult animals and embryos of *Parasteatoda tepidariorum* were obtained from the colony at UW-Madison, US.

### Transcriptome assembly for *Charinus* whip spiders

RNAlater-fixed embryos were transferred to 1.5mL tubes filled with TRIZOL (Invitrogen) after two months, and subject to RNA extraction. Total RNA extracted from each sample of the embryos of *C. ioannicitus* (three samples) and *C. israelensis* (two samples) (SI Appendix, Table S1) was submitted for library preparation at the Biotechnology Center of the University of Wisconsin-Madison. Each sample was sequenced in triplicate in an Illumina High-Seq platform using paired-end 100 bp-long read strategy at the same facility. Read quality was assessed with FastQC (Babraham Bioinformatics). Paired-end reads for *C. ioanniticus* (ISR021) and *C. israelensis* (ISR051) were compiled and *de novo* assembled using Trinity v.3.3 (Grabherr et al., 2011) enabling Trimmomatic v.0.36 to remove adapters and low-quality reads (Bolger, Lohse, & Usadel, 2014). Transcriptome quality was assessed with the Trinity package script ‘*TrinityStats.pl*’ and BUSCO v.3 (Waterhouse et al., 2017). For BUSCO, we used the ‘Arthropoda’ database and analyzed the transcriptomes filtered for the longest isoform per Trinity gene.

### RNA sequencing for differential gene expression

The total RNA extraction of each sample of *C. ioanniticu*s and *C. israelensis* embryos was sequenced in triplicate in an Illumina High-Seq platform using a single-end 100 bp-long read strategy in the same facility as described above. For *C. ioanniticus* (normal-eyes), we sequenced two biological replicates of embryos at an early embryonic stage, before eye-spot formation (ISR021-2, ISR021-3), and one sample of late embryos, after eye-spot formation (ISR021-4); For *C. israelensis* (reduced-eyes), we sequenced embryos at an early embryonic stage (ISR051-6; ISR051-4) comparable to the early stage in *C. ioanniticus* (ISR021-2, ISR021-3), as inferred by the elongated lateral profile of the body and marked furrows on the opisthosomal segments (SI Appendix, Fig. S1).

### Differential gene expression analysis in *Charinus* and identification of eye gene orthologs

Orthologs of *Drosophila melanogaster eyeless* and *twin of eyeless* (*Pax6A, Pax6B*), *sine oculis* (*soA, soB*), *orthodenticle* (*otdA, otdB*) *Optix* (*Six3.1, Six3.2*), *dachshund* (*dacA, dacB*), and *eyes absent* (*eya*) had been previously isolated in *Parasteatoda tepidariorum* (Schomburg et al., 2015, and references therein). We used as reference sequences the complete predicted transcripts for these genes from *P. tepidariorum* genome (Schwager et al., 2017), *Cupiennius salei* (Samadi et al., 2015) (for *atonal* and *Pax6*), and *D. melanogaster*, including also *atonal* and *eyegone* from the latter species. The sequences were aligned with MAFFT (v7.407) (Katoh & Standley, 2013) and the resulting alignment used to build hidden Markov model profiles for each gene (hmmbuild, from the hmmer suite v.3.3) (Finn et al., 2015). Matches to these profiles were found using hmmsearch in the reference transcriptomes of *C. ioanniticus* and *C. israelensis* as well as in the genomes of representative arthropods including *D. melanogaster* (GCA 000001215.4), *Tribolium castaneum* (GCA 000002335.3), *Daphnia magna* (GCA 003990815.1), *Strigamia maritima* (GCA 000239455.1), *Dinothrombium tinctorium* (GCA 003675995.1), *Ixodes scapularis* (GCA 002892825.2), *Tetranychus urticae* (GCA 000239435.1), *Limulus polyphemus* (GCA 000517525.1), *Tachypleus tridentatus* (GCA 004210375.1), *Centruroides sculpturatus* (GCA 000671375.2), *Parasteatoda tepidariorum* (GCA 000365465.2) and *Trichonephila clavipes* (GCA 002102615.1). These species were selected from a pool relatively recent genome assembly resources and well curated reference genomes.

Homologous sequences (those with hmmer expectation value, e < 10^10^) to the genes of interest were then compiled into individual gene FASTA files, combined with the reference sequences used for the homology search, aligned (MAFFT), trimmed of gap rich regions (trimAL v.1.2, –gappyout) (Capella-Gutiérrez, Silla-Martínez, & Gabaldón, 2009) and used for maximum likelihood gene tree estimation (IQTREE v.1.6.8, –mset LG,WAG,JTT,DCMUT –bb 1000) (Nguyen, Schmidt, Haeseler, & Minh, 2015). The association of transcripts in the *Charinus* species with the genes of interest is based on the gene phylogeny and was followed by inspection of the coding sequences to distinguish splicing variants from other gene copies. Alignments and the list of *Charinus* sequences is available in SI Appendix Dataset S1. These gene transcript association was then used for the transcript to gene map required for the DGE analysis.

### Read mapping, transcript abundance quantification

For the *in-silico* analysis of gene expression, single-end raw reads were first trimmed using the software Trimmomatic v. 0.35 (Bolger et al., 2014). For the intra-specific analysis of early (before eyespot) and late (eyespot) embryos of *C. ioanniticus* (Comparison 1), the trimmed reads were quantified in the embryonic transcriptome of *C. ioanniticus*. For the intra-specific comparison of early embryos of *C. ioanniticus* and *C. israelensis*, two reciprocal analysis were conducted: reads from both species mapped onto *C. israelensis* transcriptome as the reference (Comparison 2.1); and reads from both species mapped onto *C. ioanniticus* transcriptome (Comparison 2.2).

Transcript abundance was quantified using the software Salmon v. 1.1.0 (Patro et al., 2017), enabling ‘*–validateMapping’* flag. Analysis of differential gene expression was conducted with the software DESeq2 v 1.24.0 (Love et al., 2014) following a pipeline with the R package *tximport* v.1.12.3 (Soneson, Love, & Robinson, 2015). The exact procedures are documented in the custom R script (SI Appendix, Dataset S2)

### Parental RNA interference, in situ hybridization, and imaging in *Parasteatoda tepidariorum*

Total RNA from a range of embryonic stages of *P. tepidariorum* was extracted with TRIZOL (Invitrogen), and cDNA was synthetized using SuperScriptIII (Invitrogen). Gene fragments for *Ptep-sine oculis A* (*soA*), *orthodenticle B* (*otdB*) and *OptixB* were amplified from cDNA using gene specific primers designed with Primers3Web version 4.1.0 (Koressaar & Remm, 2007) and appended with T7 ends. Cloning amplicons were generated using the TOPO TA Cloning Kit with One Shot Top10 chemically competent *Escherichia coli* (Invitrogen). Amplicon identities and directionality were assessed with Sanger sequencing. Primer, amplicon sequences and fragment lengths are available in SI Appendix Dataset S3. Double-stranded RNA for *Ptep-soA, Ptep-otdB* and *Ptep-OptixB* was synthetized from amplicon on plasmids using MEGAScript T7 transcription kit (Thermo Fischer) with T7/T7T3 primers. Sense and antisense RNA probes for colorimetric in situ hybridization were synthetized from plasmid templates with DIG RNA labeling mix (Roche) and T7/T3 RNA polymerase (New England Biolabs) using the manufacturer’s instructions.

Parental RNA interference (RNAi) followed established protocols for double-stranded RNA (dsRNA) injection in virgin females of *P. tepidariorum* (Oda & Akiyama-Oda, 2020). Each female was injected four times with 2.5 μL of dsRNA at a concentration of 2 μg/uL, to a total of 20μg. For *Ptep-soA*, seven virgin females were injected with dsRNA of a 1048bp cloned fragment (SI Appendix, Fig. S13C) and 3 females were injected with the same volume of dH_2_O as a procedural control. Two virgin females were injected with dsRNA for *Ptep-otdB*, and six females for *Ptep-OptixB*. All females were mated after the second injection, and were fed approximately every-other day after the last injection. Cocoons were collected until the sixth clutch, approximately one per week.

Hatchlings for all cocoons were fixed between 24–48 hours after hatching. Freshly hatched postembryos have almost no external signs of eye lenses and pigments. The selected fixation window encompasses a period in which postembryos have deposited eye pigments until the beginning of the first instar, where eyes are completely formed (SI Appendix, Video S1, S2). Hatchlings were immersed in 25% ethanol/PBST and stored at 4°C. For the *Ptep-soA* RNAi experiment, hatchlings were scored in four classes: (1) wild type, where all eyes were present and bilaterally symmetrical; (2) Eyes defective, where one or more eyes were reduced in size or completely absent; (3) dead/arrested; (4) Undetermined, where embryos were damaged or clearly freshly hatched. A subset of *Ptep-soA* dsRNA-injected embryos from four clutches (n=48) and of three control clutches (n=48) were further inspected in detail to assess the effects on individual eye types. Given that there is a spectrum on the intensity of pigment deposition in the medium eyes (ME), and small asymmetries on the shape of the early developing tapetum of the lateral eyes (LE) in control embryos, the following conservative criteria was adopted: ME were considered affected when asymmetry in pigmentation or lens size was detected. Both ME were only scored as affected when they were both completely missing, in order to rule out embryos were simply delayed in pigment deposition; LE were considered defective only when the tapetum was completely absent (SI Appendix, Fig. S12). Therefore, our coding does not allow detection of a phenotype consisting of delayed pigmentation.

For in situ hybridization, a subset of *Ptep-soA* dsRNA-injected embryos at stage 13/14 (Mittmann & Wolff, 2012) was fixed in a phase of heptane and 4% formaldehyde for 12–24 hours, washed in PBST, gradually dehydrated in methanol and stored at −20°C for at least 3 days before downstream procedures, after a modified protocol of Akiyama-Oda and Oda (2003). In situ hybridization followed the protocol of Akiyama-Oda and Oda (2003).

Embryos from in situ hybridization were stained with Hoechst nuclear staining and imaged in a Nikon SMZ25 fluorescence stereomicroscope mounted with a DS-Fi2 digital color camera (Nikon Elements software). For postembryos, the prosoma was dissected with fine forceps, gradually immersed in 70% Glycerol/PBS-T and mounted on glass slides. Postembryos were imaged using an Olympus DP70 color camera mounted on an Olympus BX60 epifluorescence compound microscope.

## Acknowledgements

Microscopy was performed at the Newcomb Imaging Center, Department of Botany, University of Wisconsin-Madison. Sequencing was performed at the UW-Madison Biotechnology Center. Access to computing nodes for intensive tasks was provided by the Center for High Throughput Computing (CHTC) and the Bioinformatics Resource Center (BRC) of the University of Wisconsin-Madison. Specimens were collected under permit 2018/42037, issued by the Israel National Parks Authority to E.G.R. Fieldwork in Israel was supported by a National Geographic Society Expeditions Council grant no. NGS-271R-18 to J.A.B. This work was supported by National Science Foundation (grant no. IOS-1552610) to P.P.S.

## References

Akiyama-Oda, Y., & Oda, H. (2006). Axis specification in the spider embryo: *dpp* is required for radial-to-axial symmetry transformation and *sog* for ventral patterning. Development, 133(12), 2347–2357. http://doi.org/10.1242/dev.02400

Arendt, D., Tessmar, K., Medeiros de Campos-Baptista, M. I., Dorresteijn, A., & Wittbrodt, J. (2002). Development of pigment-cup eyes in the polychaete *Platynereis dumerilii* and evolutionary conservation of larval eyes in bilateria. Development, 129(5), 1143–1154.

Ballesteros, J. A., & Sharma, P. P. (2019). A Critical Appraisal of the Placement of Xiphosura (Chelicerata) with Account of Known Sources of Phylogenetic Error. Systematic Biology. http://doi.org/10.1093/sysbio/syz011

Ballesteros, J. A., Santibáñez López, C. E., Kováč, Ĺ., Gavish-Regev, E., & Sharma, P. P. (2019). Ordered phylogenomic subsampling enables diagnosis of systematic errors in the placement of the enigmatic arachnid order Palpigradi. Proceedings. Biological Sciences, 286(1917), 20192426. http://doi.org/10.1098/rspb.2019.2426

Barnett, A. A., & Thomas, R. H. (2012). The delineation of the fourth walking leg segment is temporally linked to posterior segmentation in the mite *Archegozetes longisetosus* (Acari: Oribatida, Trhypochthoniidae). Evolution & Development, 14(4), 383–392. http://doi.org/10.1111/j.1525-142X.2012.00556.x

Barnett, A. A., & Thomas, R. H. (2013a). Posterior Hox gene reduction in an arthropod: *Ultrabithorax* and *Abdominal-B* are expressed in a single segment in the mite *Archegozetes longisetosus*. EvoDevo, 4(1), 23. http://doi.org/10.1186/2041-9139-4-23

Barnett, A. A., & Thomas, R. H. (2013b). The expression of limb gap genes in the mite *Archegozetes longisetosus* reveals differential patterning mechanisms in chelicerates. Evolution & Development, 15(4), 280–292. http://doi.org/10.1111/ede.12038

Bebenek, I. G., Gates, R. D., Morris, J., Hartenstein, V., & Jacobs, D. K. (2004). *sine oculis* in basal Metazoa. Development Genes and Evolution, 214(7), 342–351. http://doi.org/10.1007/s00427-004-0407-3

Bolger, A. M., Lohse, M., & Usadel, B. (2014). Trimmomatic: a flexible trimmer for Illumina sequence data. Bioinformatics, 30(15), 2114–2120. http://doi.org/10.1093/bioinformatics/btu170

Bradic, M., Teotónio, H., & Borowsky, R. L. (2013). The Population Genomics of Repeated Evolution in the Blind Cavefish *Astyanax mexicanus*. Molecular Biology and Evolution, 50(11), 2383–2400. http://doi.org/10.1093/molbev/mst136

Byrne, M., Koop, D., Morris, V. B., Chui, J., Wray, G. A., & Cisternas, P. (2017). Expression of genes and proteins of the Pax-Six-Eya-Dach network in the metamorphic sea urchin: Insights into development of the enigmatic echinoderm body plan and sensory structures. Developmental Dynamics, 247(1), 239–249. http://doi.org/10.1002/dvdy.24584

Cagan, R. (2009). Chapter 5-Principles of *Drosophila* Eye Differentiation. In Current Topics in Developmental Biology (Vol. 89, pp. 115–135). Elsevier. http://doi.org/10.1016/S0070-2153(09)89005-4

Capella-Gutiérrez, S., Silla-Martínez, J. M., & Gabaldón, T. (2009). trimAl: a tool for automated alignment trimming in large-scale phylogenetic analyses. Bioinformatics, 25(15), 1972–1973. http://doi.org/10.1093/bioinformatics/btp348

Carroll, S. B. (2008). Evo-Devo and an Expanding Evolutionary Synthesis: A Genetic Theory of Morphological Evolution. Cell, 134(1), 25–36. http://doi.org/10.1016/j.cell.2008.06.030

Coghill, L. M., Darrin Hulsey, C., Chaves-Campos, J., García de Leon, F. J., & Johnson, S. G. (2014). Next generation phylogeography of cave and surface *Astyanax mexicanus*. Molecular Phylogenetics and Evolution, 79, 368–374. http://doi.org/10.1016/j.ympev.2014.06.029

Cruz-López, J. A., Proud, D. N., & Pérez-González, A. (2016). When troglomorphism dupes taxonomists: morphology and molecules reveal the first pyramidopid harvestman (Arachnida, Opiliones, Pyramidopidae) from the New World. Zoological Journal of the Linnean Society, 177(3), 602–620. http://doi.org/10.1111/zoj.12382

Daly, I. M., How, M. J., Partridge, J. C., & Roberts, N. W. (2018). Complex gaze stabilization in mantis shrimp. Proceedings of the Royal Society B: Biological Sciences, 285(1878). http://doi.org/10.1098/rspb.2018.0594

Derkarabetian, S., Steinmann, D. B., & Hedin, M. (2010). Repeated and time-correlated morphological convergence in cave-dwelling harvestmen (Opiliones, Laniatores) from montane Western North America. PLoS ONE, 5(5), e10388. http://doi.org/10.1371/journal.pone.0010388

Esposito, L. A., Bloom, T., Caicedo-Quiroga, L., Alicea-Serrano, A. M., Sánchez-Ruíz, J. A., May-Collado, L. J., et al. (2015). Islands within islands: Diversification of tailless whip spiders (Amblypygi, *Phrynus*) in Caribbean caves. Molecular Phylogenetics and Evolution, 93(C), 107–117. http://doi.org/10.1016/j.ympev.2015.07.005

Finn, R. D., Clements, J., Arndt, W., Miller, B. L., Wheeler, T. J., Schreiber, F., et al. (2015). HMMER web server: 2015 update. Nucleic Acids Research, 43(W1), W30–8. http://doi.org/10.1093/nar/gkv397

Foelix, R. (2011). Biology of Spiders (Third Edition). Oxford: Oxford University Press.

Garwood, R. J., Sharma, P. P., Dunlop, J. A., & Giribet, G. (2014). A Paleozoic Stem Group to Mite Harvestmen Revealed through Integration of Phylogenetics and Development. Current Biology, 24(9), 1–7. http://doi.org/10.1016/j.cub.2014.03.039

Gehring, W. J., & Ikeo, K. (1999). *Pax 6:* mastering eye morphogenesis and eye evolution. Trends in Genetics: TIG, 15(9), 371–377. http://doi.org/10.1016/s0168-9525(99)01776-x

Giribet, G. (2018). Current views on chelicerate phylogeny—A tribute to Peter Weygoldt. Zoologischer Anzeiger - a Journal of Comparative Zoology, 273, 7–13. http://doi.org/10.1016/j.jcz.2018.01.004

Grabherr, M. G., Haas, B. J., Yassour, M., Levin, J. Z., Thompson, D. A., Amit, I., et al. (2011). Full-length transcriptome assembly from RNA-Seq data without a reference genome. Nature Biotechnology, 29(7), 644–652. http://doi.org/10.1038/nbt.1883

Grbić, M., Khila, A., Lee, K.-Z., Bjelica, A., Grbić, V., Whistlecraft, J., et al. (2007). Mity model: *Tetranychus urticae*, a candidate for chelicerate model organism. BioEssays, 29(5), 489–496. http://doi.org/10.1002/bies.20564

Grbić, M., Van Leeuwen, T., Clark, R. M., Rombauts, S., Rouzé, P., Grbić, V., et al. (2011). The genome of *Tetranychus urticae* reveals herbivorous pest adaptations. Nature, 479(7374), 487–492. http://doi.org/10.1038/nature10640

Gulia-Nuss, M., Nuss, A. B., Meyer, J. M., Sonenshine, D. E., Roe, R. M., Waterhouse, R. M., et al. (2016). Genomic insights into the *Ixodes scapularis* tick vector of Lyme disease. Nature Communications, 7(1), 10507–13. http://doi.org/10.1038/ncomms10507

Harland, D. P., Li, D., & Jackson, R. R. (2012). How jumping spiders see the world. http://doi.org/10.1093/acprof:oso/9780195334654.003.0010

Harvey, M. S. (2002). The neglected cousins: What do we know about the smaller arachnid orders? Journal of Arachnology, 30(2), 357–372. http://doi.org/10.1636/0161-8202(2002)030[0357:TNCWDW]2.0.CO;2

Harvey, M. S. (2007). The smaller arachnid orders: Diversity, descriptions and distributions from Linnaeus to the present (1758 to 2007). Zootaxa, 1668(1668), 363–380. http://doi.org/10.11646/zootaxa.1668.1.19

Hedin, M., & Thomas, S. M. (2010). Molecular systematics of eastern North American Phalangodidae (Arachnida: Opiliones: Laniatores), demonstrating convergent morphological evolution in caves. Molecular Phylogenetics and Evolution, 54(1), 107–121. http://doi.org/10.1016/j.ympev.2009.08.020

Herman, A., Brandvain, Y., Weagley, J., Jeffery, W. R., Keene, A. C., Kono, T. J. Y., et al. (2018). The role of gene flow in rapid and repeated evolution of cave-related traits in Mexican tetra, *Astyanax mexicanus*. Molecular Ecology, 27(22), 4397–4416. http://doi.org/10.1111/mec.14877

Homann, H. (1971). Die Augen der Araneae. Zeitschrift Für Morphologie Der Tiere, 69(3), 201–272. http://doi.org/10.1007/BF00277623

Howarth, F. G. (1993). High-stress subterranean habitats and evolutionary change in cave-inhabiting arthropods. The American Naturalist, 142 Suppl 1(Suppl.), S65–77. http://doi.org/10.1086/285523

Hoy, M. A., Waterhouse, R. M., Wu, K., Estep, A. S., Ioannidis, P., Palmer, W. J., et al. (2016). Genome Sequencing of the Phytoseiid Predatory Mite *Metaseiulus occidentalis* Reveals Completely Atomized Hox Genes and Superdynamic Intron Evolution. Genome Biology and Evolution, 8(6), 1762–1775. http://doi.org/10.1093/gbe/evw048

Jeffery, W. R. (2009). Chapter 8. Evolution and development in the cavefish *Astyanax*. Current Topics in Developmental Biology, 86, 191–221. http://doi.org/10.1016/S0070-2153(09)01008-4

Jemec, A., Škufca, D., Prevorčnik, S., Fišer, Ž., & Zidar, P. (2017). Comparative study of acetylcholinesterase and glutathione S-transferase activities of closely related cave and surface *Asellus aquaticus* (Isopoda: Crustacea). PLoS ONE, 12(5), e0176746–14. http://doi.org/10.1371/journal.pone.0176746

Juan, C., Guzik, M. T., Jaume, D., & Cooper, S. J. B. (2010). Evolution in caves: Darwin’s ‘wrecks of ancient life’ in the molecular era. Molecular Ecology, 19(18), 3865–3880. http://doi.org/10.1111/j.1365-294X.2010.04759.x

Katoh, K., & Standley, D. M. (2013). MAFFT multiple sequence alignment software version 7: improvements in performance and usability. Molecular Biology and Evolution, 30(4), 772–780. http://doi.org/10.1093/molbev/mst010

Kenny, N. J., Chan, K. W., Nong, W., Qu, Z., Maeso, I., Yip, H. Y., et al. (2015). Ancestral whole-genome duplication in the marine chelicerate horseshoe crabs. Heredity, 116(2, 190–199. http://doi.org/10.1038/hdy.2015.89

Khadjeh, S., Turetzek, N., Pechmann, M., Schwager, E. E., Wimmer, E. A., Damen, W. G. M., & Prpic, N. M. (2012). Divergent role of the Hox gene *Antennapedia* in spiders is responsible for the convergent evolution of abdominal limb repression. Proceedings of the National Academy of Sciences, 109(13), 4921–4926. http://doi.org/10.1073/pnas.1116421109

Koressaar, T., & Remm, M. (2007). Enhancements and modifications of primer design program Primer3. Bioinformatics, 23(10), 1289–1291. http://doi.org/10.1093/bioinformatics/btm091

Kumar, J. P. (2009). The molecular circuitry governing retinal determination. Biochimica Et Biophysica Acta (BBA) - Gene Regulatory Mechanisms, 1789(4), 306–314. http://doi.org/10.1016/j.bbagrm.2008.10.001

Land, M. F. (1985). The Morphology and Optics of Spider Eyes. In Neurobiology of Arachnids (Vol. 189, pp. 53–78). Berlin, Heidelberg: Springer, Berlin, Heidelberg. http://doi.org/10.1007/978-3-642-70348-5_4

Leite, D. J., Baudouin-Gonzalez, L., Iwasaki-Yokozawa, S., Lozano-Fernandez, J., Turetzek, N., Akiyama-Oda, Y., et al. (2018). Homeobox Gene Duplication and Divergence in Arachnids. Molecular Biology and Evolution, 35(9), 2240–2253. http://doi.org/10.1093/molbev/msy125

Love, M. I., Huber, W., & Anders, S. (2014). Moderated estimation of fold change and dispersion for RNA-seq data with DESeq2. Genome Biology, 15(12), 31–21. http://doi.org/10.1186/s13059-014-0550-8

Lozano-Fernandez, J., Tanner, A. R., Giacomelli, M., Carton, R., Vinther, J., Edgecombe, G. D., & Pisani, D. (2019). Increasing species sampling in chelicerate genomic-scale datasets provides support for monophyly of Acari and Arachnida. Nature Communications, 1–8. http://doi.org/10.1038/s41467-019-10244-7

Mammola, S., & Isaia, M. (2017). Spiders in caves. Proceedings of the Royal Society B: Biological Sciences, 284(1853), 20170193–10. http://doi.org/10.1098/rspb.2017.0193

Mammola, S., Arnedo, M. A., Pantini, P., Piano, E., Chiappetta, N., & Isaia, M. (2018a). Ecological speciation in darkness? Spatial niche partitioning in sibling subterranean spiders (Araneae: Linyphiidae: Troglohyphantes). Invertebrate Systematics, 32(5), 1069–1082. http://doi.org/10.1071/IS17090

Mammola, S., Cardoso, P., Ribera, C., Pavlek, M., & Isaia, M. (2018b). A synthesis on cave-dwelling spiders in Europe. Journal of Zoological Systematics and Evolutionary Research, 56(3), 301–316. http://doi.org/10.1111/jzs.12201

Mammola, S., Mazzuca, P., Pantini, P., Isaia, M., & Arnedo, M. A. (2017). Advances in the systematics of the spider genus *Troglohyphantes* (Araneae, Linyphiidae). Systematics and Biodiversity, 15(4), 307–326. http://doi.org/10.1080/14772000.2016.1254304

Marshall, J., Cronin, T. W., & Kleinlogel, S. (2007). Stomatopod eye structure and function: A review. Arthropod Structure & Development, 36(4), 420–448. http://doi.org/10.1016/j.asd.2007.01.006

Miranda, G. S., Aharon, S., Gavish-Regev, E., Giupponi, A. P. L., & Wizen, G. (2016). A new species of *Charinus* Simon, 1892 (Arachnida: Amblypygi: Charinidae) from Israel and new records of *C. ioanniticus* (Kritscher, 1959). European Journal of Taxonomy, (234), 1–18. http://doi.org/10.5852/ejt.2016.234

Mittmann, B., & Wolff, C. (2012). Embryonic development and staging of the cobweb spider *Parasteatoda tepidariorum* C. L. Koch, 1841 (syn.: *Achaearanea tepidariorum;* Araneomorphae; Theridiidae). Development Genes and Evolution, 222(4), 189–216. http://doi.org/10.1007/s00427-012-0401-0

Mojaddidi, H., Fernandez, F. E., Erickson, P. A., & Protas, M. E. (2018). Embryonic origin and genetic basis of cave associated phenotypes in the isopod crustacean *Asellus aquaticus*. Scientific Reports, 1–12. http://doi.org/10.1038/s41598-018-34405-8

Morehouse, N. I., Buschbeck, E. K., Zurek, D. B., Steck, M., & Porter, M. L. (2017). Molecular Evolution of Spider Vision: New Opportunities, Familiar Players. Biol Bull, 233(1), 21–38. http://doi.org/10.1086/693977

Nguyen, L.-T., Schmidt, H. A., Haeseler, von, A., & Minh, B. Q. (2015). IQ-TREE: a fast and effective stochastic algorithm for estimating maximum-likelihood phylogenies. Molecular Biology and Evolution, 32(1), 268–274. http://doi.org/10.1093/molbev/msu300

Nolan, E. D., Santibáñez López, C. E., & Sharma, P. P. (2020). Developmental gene expression as a phylogenetic data class: support for the monophyly of Arachnopulmonata. Development Genes and Evolution, 230(2), 137–153. http://doi.org/10.1007/s00427-019-00644-6

Nossa, C. W., Havlak, P., Yue, J.-X., Lv, J., Vincent, K. Y., Brockmann, H. J., & Putnam, N. H. (2014). Joint assembly and genetic mapping of the Atlantic horseshoe crab genome reveals ancient whole genome duplication. GigaScience, 3(1), 708–21. http://doi.org/10.1186/2047-217X-3-9

Oda, H., & Akiyama-Oda, Y. (2020). The common house spider *Parasteatoda tepidariorum*. EvoDevo, 1–7. http://doi.org/10.1186/s13227-020-00152-z

Paese, C. L. B., Leite, D. J., Schönauer, A., McGregor, A. P., & Russell, S. (2018). Duplication and expression of Sox genes in spiders, 1–14. http://doi.org/10.1186/s12862-018-1337-4

Patro, R., Duggal, G., Love, M. I., Irizarry, R. A., & Kingsford, C. (2017). Salmon provides fast and bias-aware quantification of transcript expression. Nature Publishing Group, 14(4), 417–419. http://doi.org/10.1038/nmeth.4197

Paulus, H. F. (1979). Eye structure and the monophyly of the Arthropoda. In “Arthropod Phylogeny”(A. PGupta, Ed.) pp. 299–383.

Pechmann, M. (2016). Formation of the germ-disc in spider embryos by a condensation-like mechanism. Frontiers in Zoology, 13(1), 35. http://doi.org/10.1186/s12983-016-0166-9

Pechmann, M., McGregor, A. P., Schwager, E. E., Feitosa, N. M., & Damen, W. G. M. (2009). Dynamic gene expression is required for anterior regionalization in a spider. Proceedings of the National Academy of Sciences of the United States of America, 106(5), 1468–1472. http://doi.org/10.1073/pnas.0811150106

Pineda, D., Gonzalez, J., Callaerts, P., Ikeo, K., Gehring, W. J., & Salo, E. (2000). Searching for the prototypic eye genetic network: *Sine oculis* is essential for eye regeneration in planarians. Proceedings of the National Academy of Sciences, 97(9), 4525–4529. http://doi.org/10.1073/pnas.97.9.4525

Porter, M. L., Dittmar, K., & Pérez-Losada, M. (2007). How long does evolution of the troglomorphic form take? Estimating divergence times in *Astyanax mexicanus*. Acta Carsologica, 36(1), 173–182. http://doi.org/10.3986/ac.v36i1.219

Posnien, N., Zeng, V., Schwager, E. E., Pechmann, M., Hilbrant, M., Keefe, J. D., et al. (2014). A Comprehensive Reference Transcriptome Resource for the Common House Spider *Parasteatoda tepidariorum*. PLoS ONE, 9(8), e104885–20. http://doi.org/10.1371/journal.pone.0104885

Protas, M. E., Hersey, C., Kochanek, D., Zhou, Y., Wilkens, H., Jeffery, W. R., et al. (2005). Genetic analysis of cavefish reveals molecular convergence in the evolution of albinism. Nature Genetics, 38(1), 107–111. http://doi.org/10.1038/ng1700

Protas, M. E., Trontelj, P., & Patel, N. H. (2011). Genetic basis of eye and pigment loss in the cave crustacean, *Asellus aquaticus*. Proceedings of the National Academy of Sciences of the United States of America, 108(14), 5702–5707. http://doi.org/10.1073/pnas.1013850108

Protas, M., & Jeffery, W. R. (2012). Evolution and development in cave animals: from fish to crustaceans. Wiley Interdisciplinary Reviews: Developmental Biology, 1(6), 823–845. http://doi.org/10.1002/wdev.61

Re, C., Fišer, Ž., Perez, J., Tacdol, A., Trontelj, P., & Protas, M. E. (2018). Common Genetic Basis of Eye and Pigment Loss in Two Distinct Cave Populations of the Isopod Crustacean *Asellus aquaticus*. Integrative and Comparative Biology, 58(3), 421–430. http://doi.org/10.1093/icb/icy028

Riddle, M. R., Aspiras, A. C., Gaudenz, K., Peuß, R., Sung, J. Y., Martineau, B., et al. (2018). Insulin resistance in cavefish as an adaptation to a nutrient-limited environment. Nature, 1–19. http://doi.org/10.1038/nature26136

Rivera, A., Winters, I., Rued, A., Ding, S., Posfai, D., Cieniewicz, B., et al. (2013). The evolution and function of the Pax/Six regulatory network in sponges. Evolution & Development, 15(3), 186–196. http://doi.org/10.1111/ede.12032

Rota-Stabelli, O., Campbell, L., Brinkmann, H., Edgecombe, G. D., Longhorn, S. J., Peterson, K. J., et al. (2010). A congruent solution to arthropod phylogeny: phylogenomics, microRNAs and morphology support monophyletic Mandibulata. Proceedings of the Royal Society B: Biological Sciences, 278(1703), 298–306. http://doi.org/10.1098/rspb.2010.0590

Samadi, L., Schmid, A., & Eriksson, B. J. (2015). Differential expression of retinal determination genes in the principal and secondary eyes of *Cupiennius salei* Keyserling (1877). EvoDevo, 6(1), 16–17. http://doi.org/10.1186/s13227-015-0010-x

Sandve, S. R., Rohlfs, R. V., & Hvidsten, T. R. (2018). Subfunctionalization versus neofunctionalization after whole-genome duplication. Nature Publishing Group, 50(7), 908–909. http://doi.org/10.1038/s41588-018-0162-4

Santibáñez López, C. E., Francke, O. F., & Prendini, L. (2014). Shining a light into the world’s deepest caves: phylogenetic systematics of the troglobiotic scorpion genus *Alacran* Francke, 1982 (Typhlochactidae:Alacraninae). Invertebrate Systematics, 28(6), 643–664. http://doi.org/10.1071/IS14035

Santos, V. T., Ribeiro, L., Fraga, A., de Barros, C. M., Campos, E., Moraes, J., et al. (2013). The embryogenesis of the Tick *Rhipicephalus* (*Boophilus*) *microplus*: The establishment of a new chelicerate model system. Genesis, 51(12), 803–818. http://doi.org/10.1002/dvg.22717

Schacht, M. I., Schomburg, C., & Bucher, G. (2020). *six3* acts upstream of *foxQ2* in labrum and neural development in the spider *Parasteatoda tepidariorum*. Development Genes and Evolution, 230(2), 95–104. http://doi.org/10.1007/s00427-020-00654-9

Schomburg, C., Turetzek, N., Schacht, M. I., Schneider, J., Kirfel, P., Prpic, N.-M., & Posnien, N. (2015). Molecular characterization and embryonic origin of the eyes in the common house spider *Parasteatoda tepidariorum*. EvoDevo, 6(1), 15. http://doi.org/10.1186/s13227-015-0011-9

Schwager, E. E., Pechmann, M., Feitosa, N. M., McGregor, A. P., & Damen, W. G. M. (2009). *hunchback* functions as a segmentation gene in the spider *Achaearanea tepidariorum*. Current Biology: CB, 19(16), 1333–1340. http://doi.org/10.1016/j.cub.2009.06.061

Schwager, E. E., Sharma, P. P., Clarke, T., Leite, D. J., Wierschin, T., Pechmann, M., et al. (2017). The house spider genome reveals an ancient whole-genome duplication during arachnid evolution. BMC Biology, 15(1), 62. http://doi.org/10.1186/s12915-017-0399-x

Setton, E. V. W., March, L. E., Nolan, E. D., Jones, T. E., Cho, H., Wheeler, W. C., et al. (2017). Expression and function of *spineless* orthologs correlate with distal deutocerebral appendage morphology across Arthropoda. Developmental Biology, 430(1), 224–236. http://doi.org/10.1016/j.ydbio.2017.07.016

Sharma, P. P., Kaluziak, S. T., Pérez-Porro, A. R., González, V. L., Hormiga, G., Wheeler, W. C., & Giribet, G. (2014a). Phylogenomic interrogation of Arachnida reveals systemic conflicts in phylogenetic signal. Molecular Biology and Evolution, 31(11), 2963–2984. http://doi.org/10.1093/molbev/msu235

Sharma, P. P., Santiago, M. A., González-Santillán, E., Monod, L., & Wheeler, W. C. (2015a). Evidence of duplicated Hox genes in the most recent common ancestor of extant scorpions. Evolution & Development, 17(6), 347–355. http://doi.org/10.1111/ede.12166

Sharma, P. P., Schwager, E. E., Extavour, C. G., & Giribet, G. (2012). Hox gene expression in the harvestman *Phalangium opilio* reveals divergent patterning of the chelicerate opisthosoma. Evolution & Development, 14(5), 450–463. http://doi.org/10.1111/j.1525-142X.2012.00565.x

Sharma, P. P., Schwager, E. E., Extavour, C. G., & Wheeler, W. C. (2014b). Hox gene duplications correlate with posterior heteronomy in scorpions. Proceedings. Biological Sciences, 281(1792), 20140661–20140661. http://doi.org/10.1098/rspb.2014.0661

Sharma, P. P., Schwager, E. E., Giribet, G., Jockusch, E. L., & Extavour, C. G. (2013). *Distal-less* and *dachshund* pattern both plesiomorphic and apomorphic structures in chelicerates: RNA interference in the harvestman *Phalangium opilio* (Opiliones). Evolution & Development, 15(4), 228–242. http://doi.org/10.1111/ede.12029

Sharma, P. P., Tarazona, O. A., Lopez, D. H., Schwager, E. E., Cohn, M. J., Wheeler, W. C., & Extavour, C. G. (2015b). A conserved genetic mechanism specifies deutocerebral appendage identity in insects and arachnids. Proceedings of the Royal Society B: Biological Sciences, 282(1808), 20150698–20150698. http://doi.org/10.1098/rspb.2015.0698

Shubin, N., Tabin, C., & Carroll, S. (2009). Deep homology and the origins of evolutionary novelty. Nature, 457(7231), 818–823. http://doi.org/10.1038/nature07891

Shull, L. C., Sen, R., Menzel, J., Goyama, S., Kurokawa, M., & Artinger, K. B. (2020). The conserved and divergent roles of Prdm3 and Prdm16 in zebrafish and mouse craniofacial development. Developmental Biology. http://doi.org/10.1016/j.ydbio.2020.02.006

Smrž, J., Kováč, Ĺ., Mikeš, J., & Lukešová, A. (2013). Microwhip Scorpions (Palpigradi) Feed on Heterotrophic Cyanobacteria in Slovak Caves - A Curiosity among Arachnida. PLoS ONE, 8(10), e75989. http://doi.org/10.1371/journal.pone.0075989

Soneson, C., Love, M. I., & Robinson, M. D. (2015). Differential analyses for RNA-seq: transcript-level estimates improve gene-level inferences. F1000Research, 4, 1521. http://doi.org/10.12688/f1000research.7563.1

Stahl, B. A., Gross, J. B., Speiser, D. I., Oakley, T. H., Patel, N. H., Gould, D. B., & Protas, M. E. (2015). A Transcriptomic Analysis of Cave, Surface, and Hybrid Isopod Crustaceans of the Species *Asellus aquaticus*. PLoS ONE, 10(10), e0140484–14. http://doi.org/10.1371/journal.pone.0140484

Strickler, A. G., Yamamoto, Y., & Jeffery, W. R. (2001). Early and late changes in *Pax6* expression accompany eye degeneration during cavefish development. Development Genes and Evolution, 211(3), 138–144. http://doi.org/10.1007/s004270000123

Takagi, A., Kurita, K., Terasawa, T., Nakamura, T., Bando, T., Moriyama, Y., et al. (2012). Functional analysis of the role of *eyes absent* and *sine oculis* in the developing eye of the cricket *Gryllus bimaculatus*. Development, Growth & Differentiation, 54(2), 227–240. http://doi.org/10.1111/j.1440-169X.2011.01325.x

Telford, M. J., & Thomas, R. H. (1998). Expression of homeobox genes shows chelicerate arthropods retain their deutocerebral segment. Proceedings of the National Academy of Sciences, 95(18), 10671–10675. http://doi.org/10.1073/pnas.95.18.10671

Thoen, H. H., How, M. J., Chiou, T.-H., & Marshall, J. (2014). A different form of color vision in mantis shrimp. Science, 343(6169), 411–413. http://doi.org/10.1126/science.1245824

Turetzek, N., Pechmann, M., Schomburg, C., Schneider, J., & Prpic, N.-M. (2015). Neofunctionalization of a Duplicate *dachshund* Gene Underlies the Evolution of a Novel Leg Segment in Arachnids. Molecular Biology and Evolution, 33(1), 109–121. http://doi.org/10.1093/molbev/msv200

Vopalensky, P., & Kozmik, Z. (2009). Eye evolution: common use and independent recruitment of genetic components. Philosophical Transactions of the Royal Society B: Biological Sciences, 364(1531), 2819–2832. http://doi.org/10.1098/rstb.2009.0079

Waterhouse, R. M., Seppey, M., Simão, F. A., Manni, M., Ioannidis, P., Klioutchnikov, G., et al. (2017). BUSCO Applications from Quality Assessments to Gene Prediction and Phylogenomics. Molecular Biology and Evolution, 35(3), 543–548. http://doi.org/10.1093/molbev/msx319

Weygoldt, P. (1975). Untersuchungen zur Embryologie und Morphologie der Geißelspinne *Tarantula marginemaculata* C. L. Koch (Arachnida, Amblypygi, Tarantulidae). Zoomorphologie, 82, 137–199.

Weygoldt, P. (2000). Whip Spiders (Chelicerata: Amblypygi). Their Biology, Morphology and Systematics. Apollo Books.

ZarinKamar, N., Yang, X., Bao, R., Friedrich, F., Beutel, R., & Friedrich, M. (2011). The *Pax* gene *eyegone* facilitates repression of eye development in *Tribolium*. EvoDevo, 2(1), 8–15. http://doi.org/10.1186/2041-9139-2-8

Zhou, Y., Liang, Y., Yan, Q., Zhang, L., Chen, D., Ruan, L., et al. (2020). The draft genome of horseshoe crab *Tachypleus tridentatus* reveals its evolutionary scenario and well-developed innate immunity. BMC Genomics, 21(1), 137–15. http://doi.org/10.1186/s12864-020-6488-1

Zurek, D. B., Cronin, T. W., Taylor, L. A., Byrne, K., Sullivan, M. L. G., & Morehouse, N. I. (2015). Spectral filtering enables trichromatic vision in colorful jumping spiders. Current Biology, 25(10), R403–R404. http://doi.org/10.1016/j.cub.2015.03.033

