## Supplementary material for "How spiders make their eyes: Systemic paralogy and function of retinal determination network homologs in arachnids": SI Appendix Figure S1-S13; Table S1-S2

*Equal author contribution

**Supplementary Information Appendix**

Table of Contents

**Table of Contents** ... 1

***SI Appendix, Figure S1*** ... 3

***SI Appendix, Figure S2:*** ... 4

***SI Appendix, Figure S3:*** ... 5

***SI Appendix, Figure S4:*** ... 6

***SI Appendix, Figure S5:*** ... 7

***SI Appendix, Figure S6:*** ... 8

***SI Appendix, Figure S7:*** ... 9

***SI Appendix, Figure S8:*** ... 10

***SI Appendix, Figure S9:*** ... 11

***SI Appendix, Figure S10:*** ... 12

***SI Appendix, Figure S11:*** ... 13

***SI Appendix, Figure S12:*** ... 15

***SI Appendix, Figure S13:*** ... 16

***SI Appendix, Video S1:*** ... 17

***SI Appendix, Video S2:*** ... 17

***SI Appendix, Dataset S1:*** ... 17

***SI Appendix, Dataset S2:*** ... 17

***SI Appendix, Dataset S3:*** ... 17

***SI Appendix, Table S1:*** ... 19

***SI Appendix, Table S2:*** ... 20

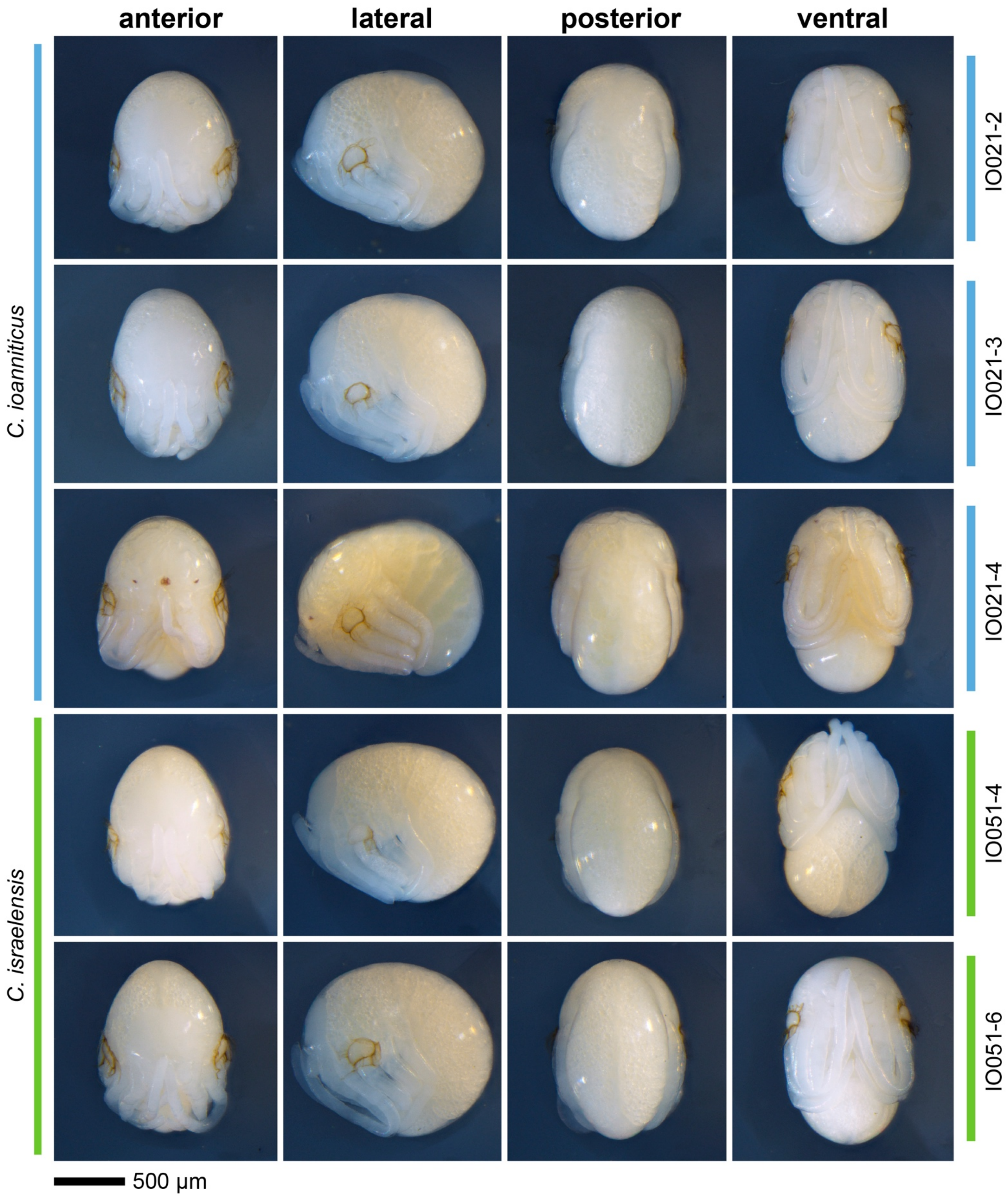


***SI Appendix, Figure S1*****:** Voucher specimens of the deutembryo stages *Charinus ioanniticus* (top three rows) and *Charinus israelensis* (top bottom rows) used in each RNA extraction. Each row is a different view of the same embryo. IO051-2 and IO051-3: early deutembryos pre-eyespots, *C. ioanniticus*. IO021-4: late deutembryos with eyespots, *C. ioanniticus*. IO051-4 and IO051-6: early deutembryos.

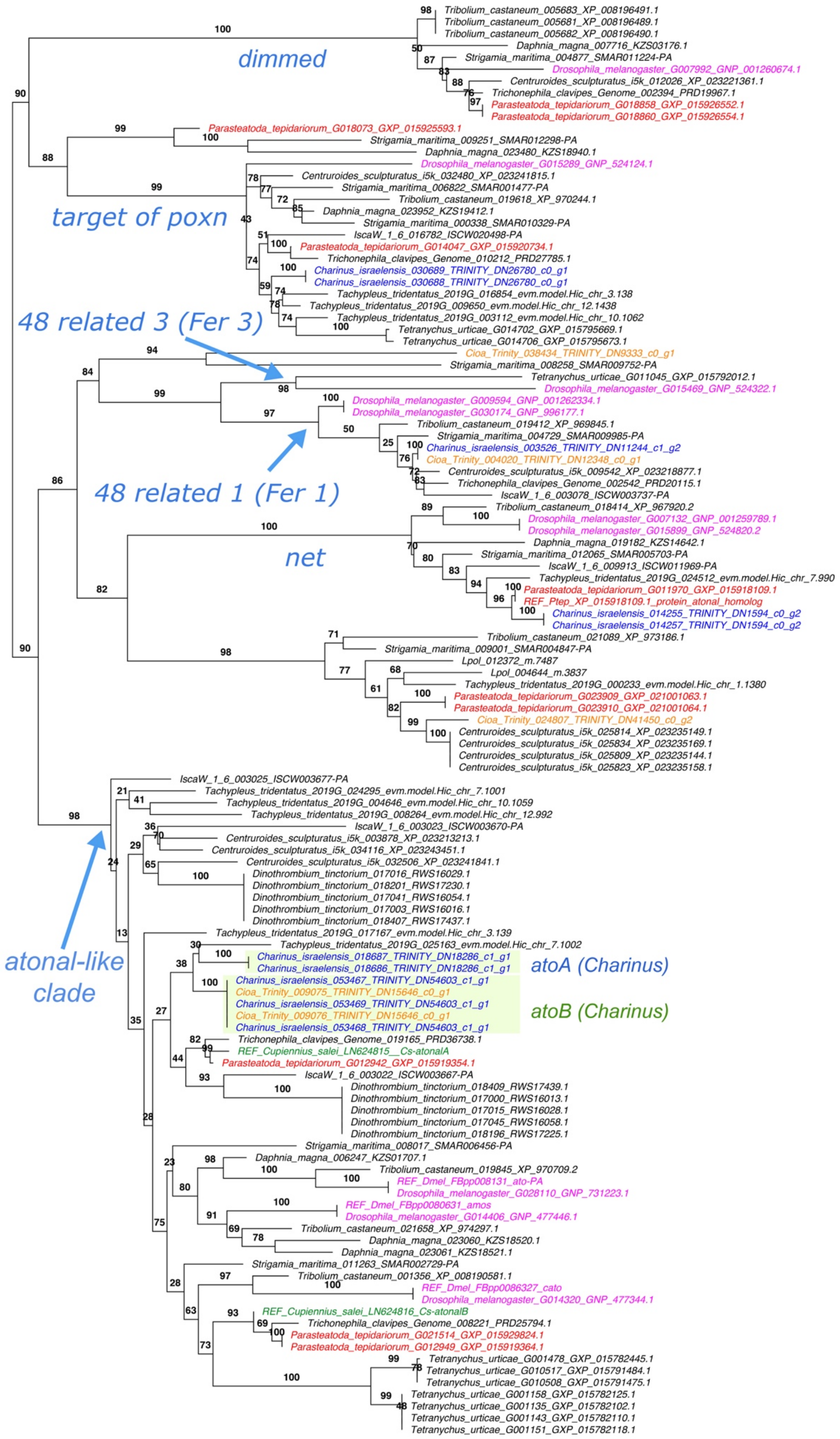


***SI Appendix, Figure S2:*** Gene tree of *atonal* homologs. Multiple isoforms per gene were included for most terminals. Reference sequences of *Drosophila melanogaster* (cyan), *Parasteatoda tepidariorum* (red), and *Cupiennius salei* (green) were used to inform annotation (see Material and Methods). Terminals for *Charinus ioanniticus* and *C. israelensis* are colored orange and blue, respectively.

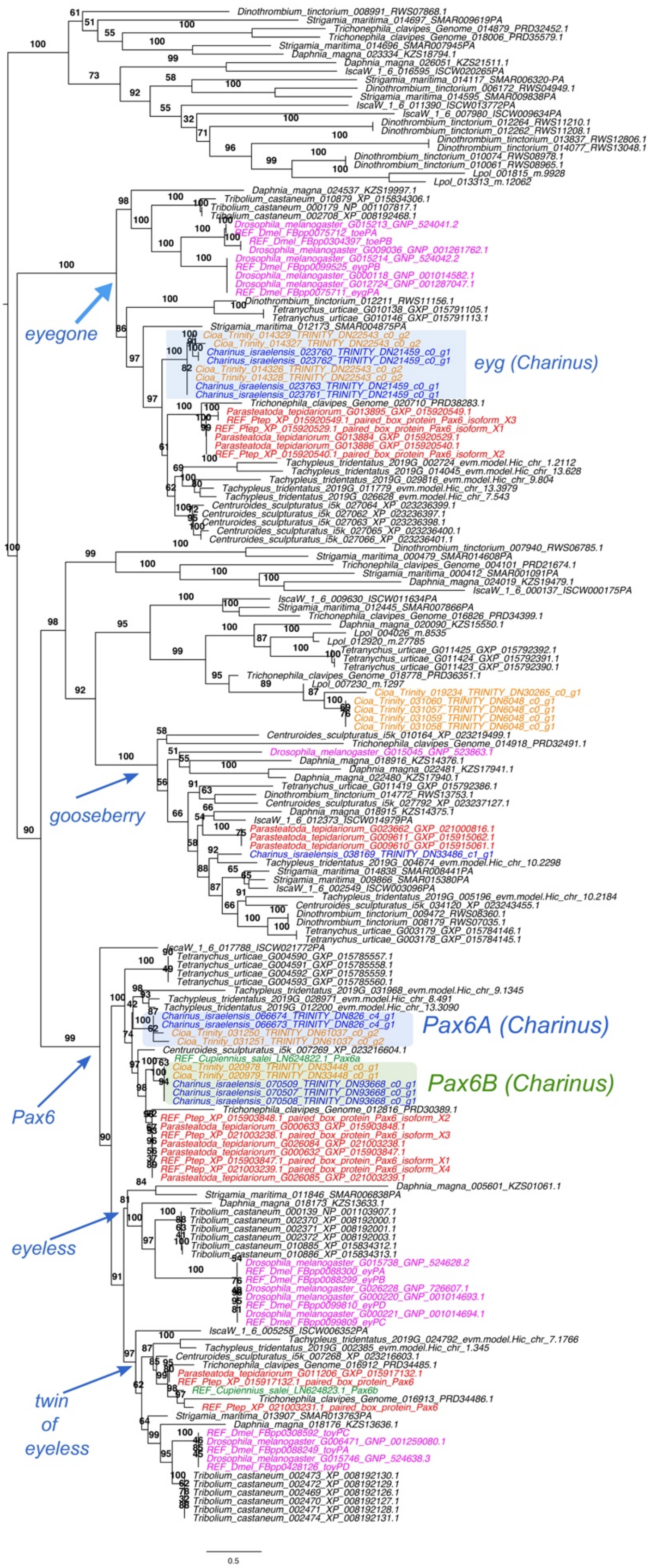


***SI Appendix, Figure S3:*** Gene tree of *Pax6* and *eyegone* homologs. Multiple isoforms per gene were included for most terminals. Reference sequences of *Drosophila melanogaster* (cyan), *Parasteatoda tepidariorum* (red), and *Cupiennius salei* (green) were used to inform annotation (see Material and Methods). Terminals for *Charinus ioanniticus* and *C. israelensis* are colored orange and blue, respectively.

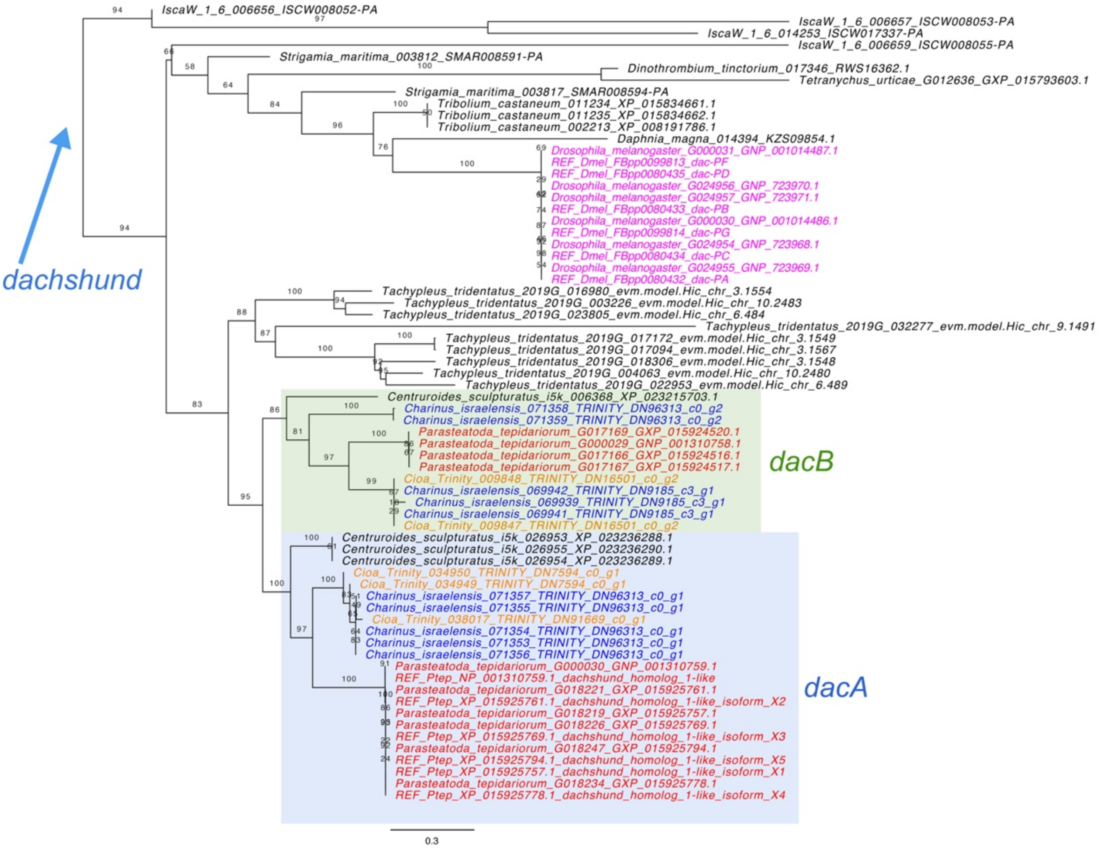


***SI Appendix, Figure S4:*** Gene tree of *dachshund* homologs. Multiple isoforms per gene were included for most terminals. Reference sequences of *Drosophila melanogaster* (cyan) and *Parasteatoda tepidariorum* (red) were used to inform annotation (see Material and Methods). Terminals for *Charinus ioanniticus* and *C. israelensis* are colored orange and blue, respectively.The two well-defined clades of *C. israelensis* sequences in the *dacB* clade overlap in only three amino acids, and are probably two fragmentary assemblies of the same gene. *Ixodes scapularis* (*Isca*) has at least four *dac* copies which appear to be specific to this species.

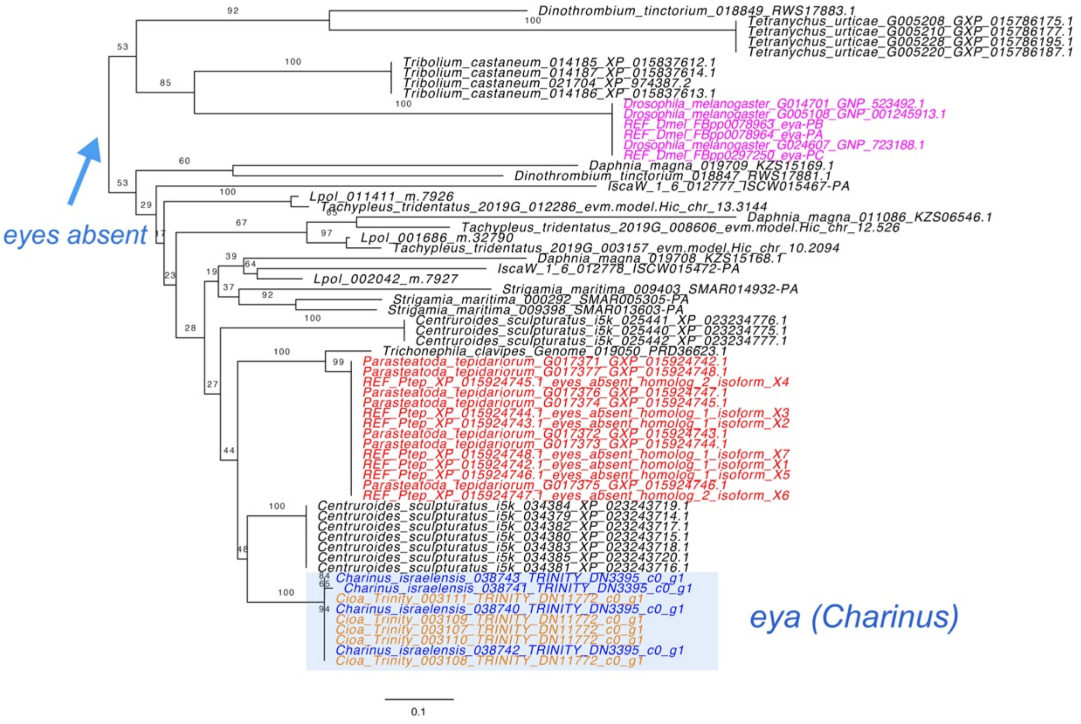


***SI Appendix, Figure S5:*** Gene tree of *eyes absent* homologs. Multiple isoforms per gene were included for most terminals. Reference sequences of *Drosophila melanogaster* (cyan) and *Parasteatoda tepidariorum* (red) were used to inform annotation (see Material and Methods). Terminals for *Charinus ioanniticus* and *C. israelensis* are colored orange and blue, respectively. The terminals for *Ixodes scapularis* (*Isca*) and *Dinothrombium tinctorium* each appear to be fragmentary assemblies of a single copy, as they have little overlapping sequence between them (see alignment in Appendix Dataset S1)

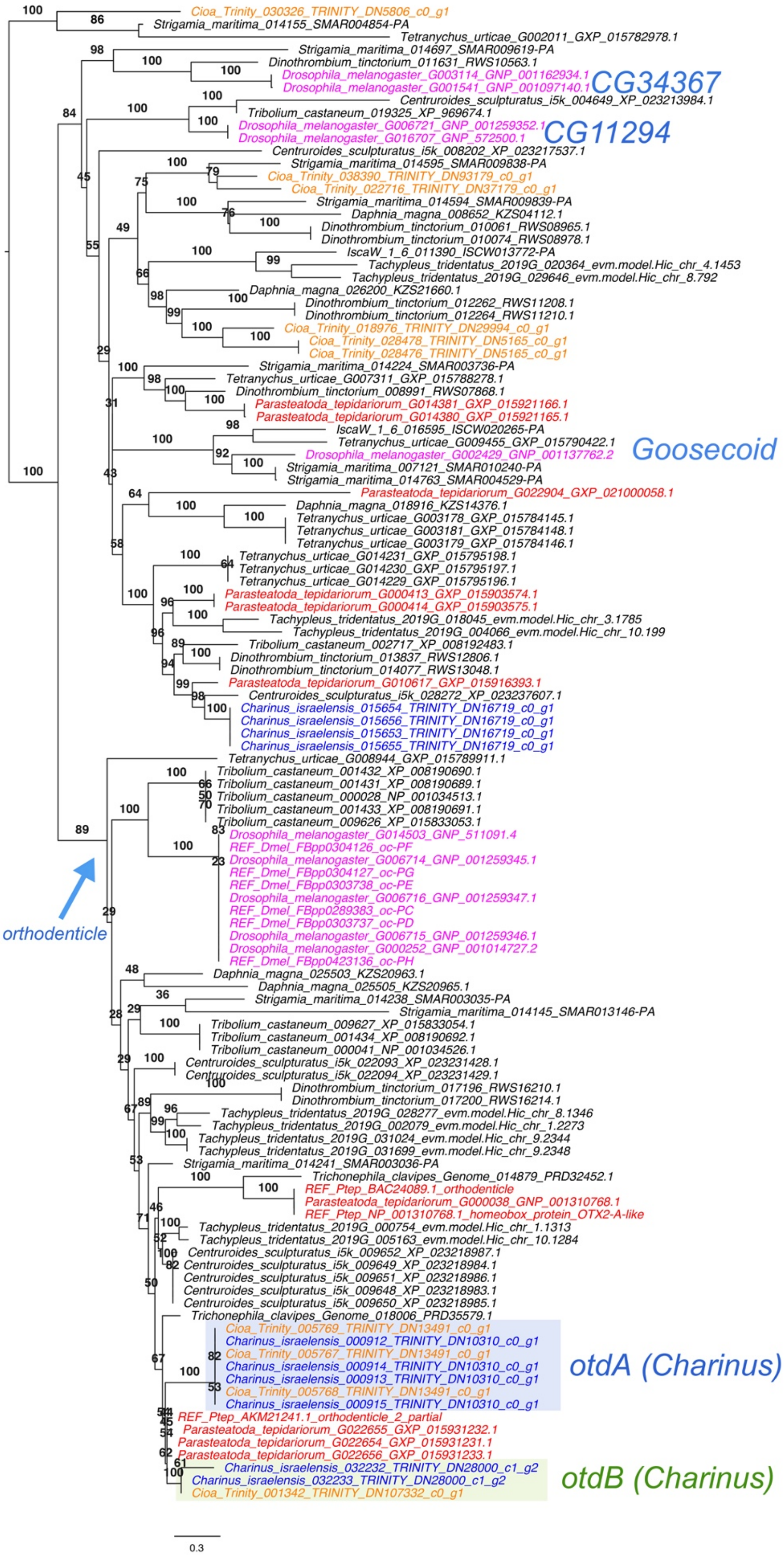


***SI Appendix, Figure S6:*** Gene tree of *goosecoid* and *orthodenticle* homologs. Multiple isoforms per gene were included for most terminals. Reference sequences of *Drosophila melanogaster* (cyan) and *Parasteatoda tepidariorum* (red) were used to inform annotation (see Material and Methods). Terminals for *Charinus ioanniticus* and *C. israelensis* are colored orange and blue, respectively.

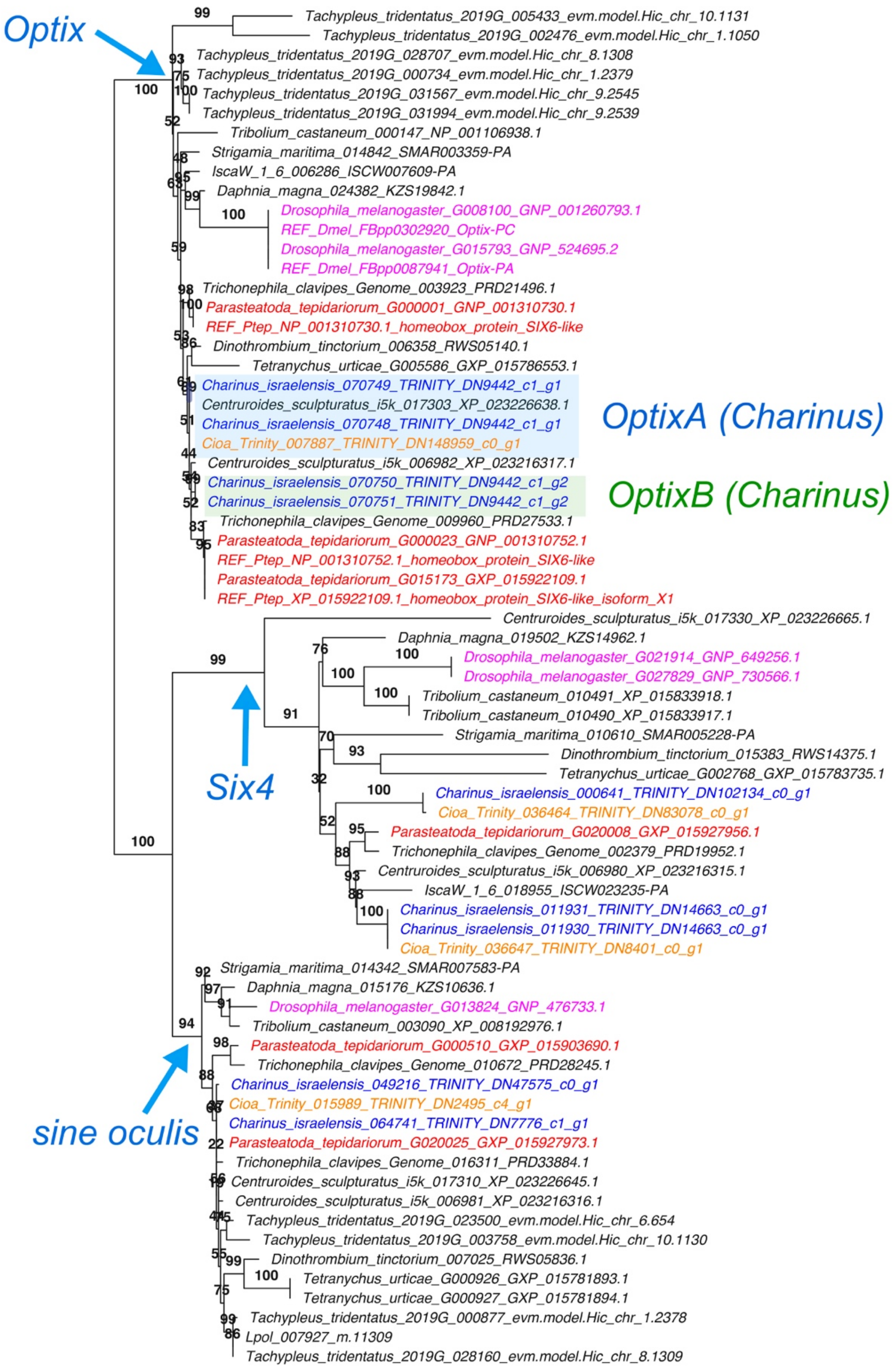


***SI Appendix, Figure S7:*** Gene tree of *Six* genes homologs, including *Six1* (*sineoculis*), *Six3* (*Optix*) and *Six4*. Multiple isoforms per gene were included for most terminals. Reference sequences of *Drosophila melanogaster* (cyan) and *Parasteatoda tepidariorum* (red) were used to inform annotation (see Material and Methods). Terminals for *Charinus ioannitiucus* and *C. israelensis* are colored orange and blue, respectively.

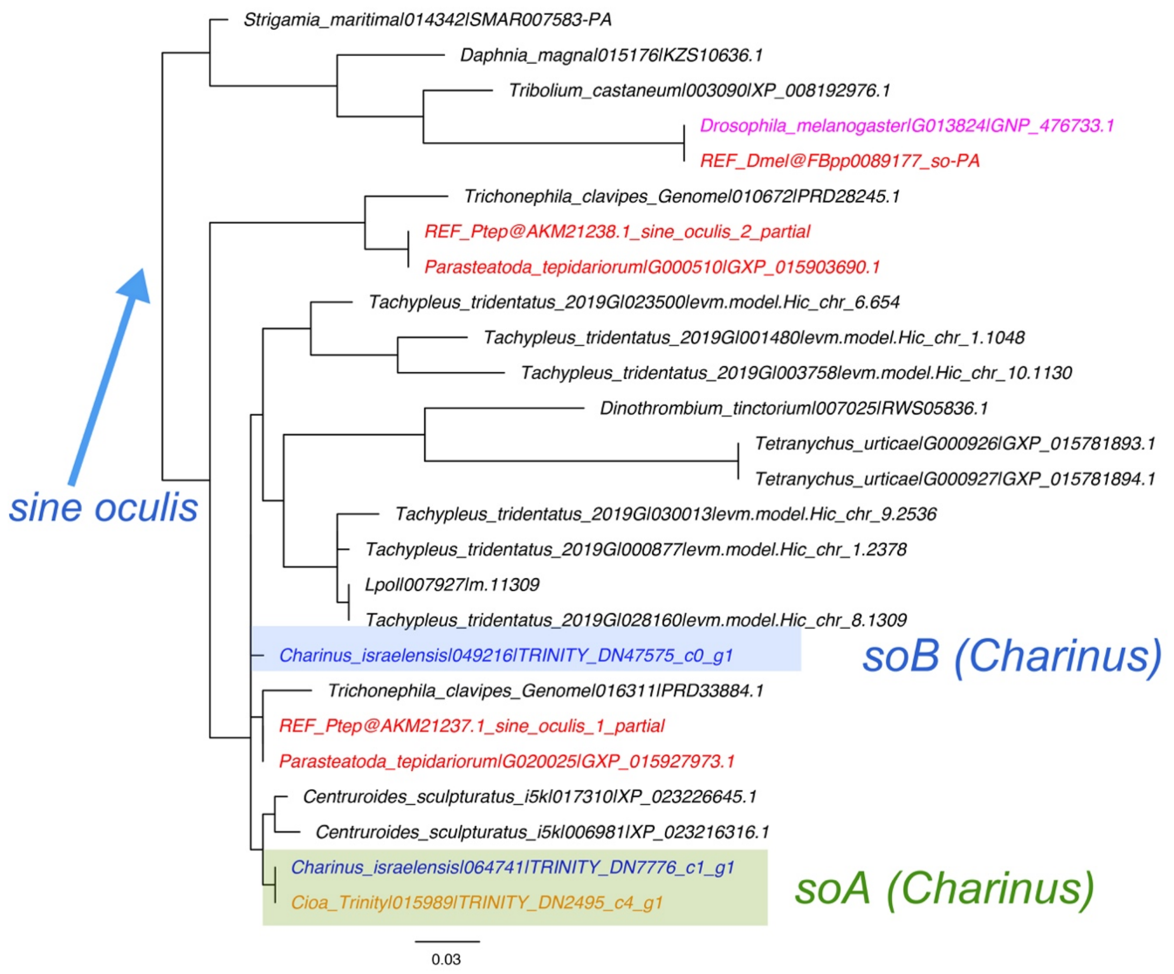


***SI Appendix, Figure S8:*** Gene tree of *sineoculis* homologs. Multiple isoforms per gene were included for most terminals. Reference sequences of *Drosophila melanogaster* (cyan) and *Parasteatoda tepidariorum* (red) were used to inform annotation (see Material and Methods). Terminals for *Charinus ioanniticus* and *C. israelensis* are colored orange and blue, respectively.

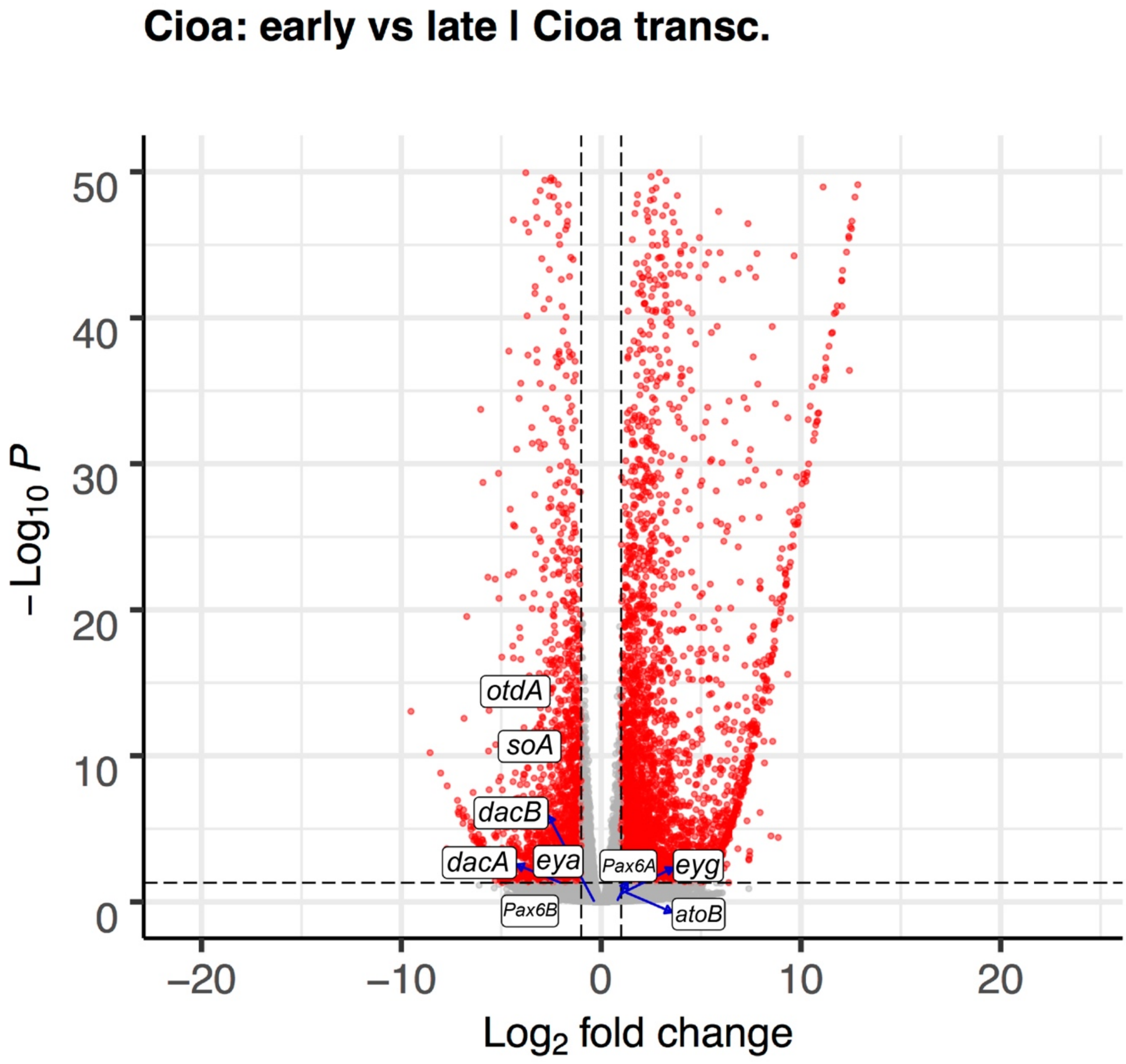


***SI Appendix, Figure S9:*** Comparison 1. Volcano plot of $p_{adj}$ values (y-axis) and $log_2$ fold change (x-axis) of all the genes in the analysis differential gene expression comparing reads from early deutembryos of *C. ioanniticus* (before eyespots) versus late deutembryos of *C. ioanniticus* (with eyespots). Each gene is represented by a dot. Red dots have $p_{adj}$>0.05. Dashed lines mark $log_2$FC > [1].

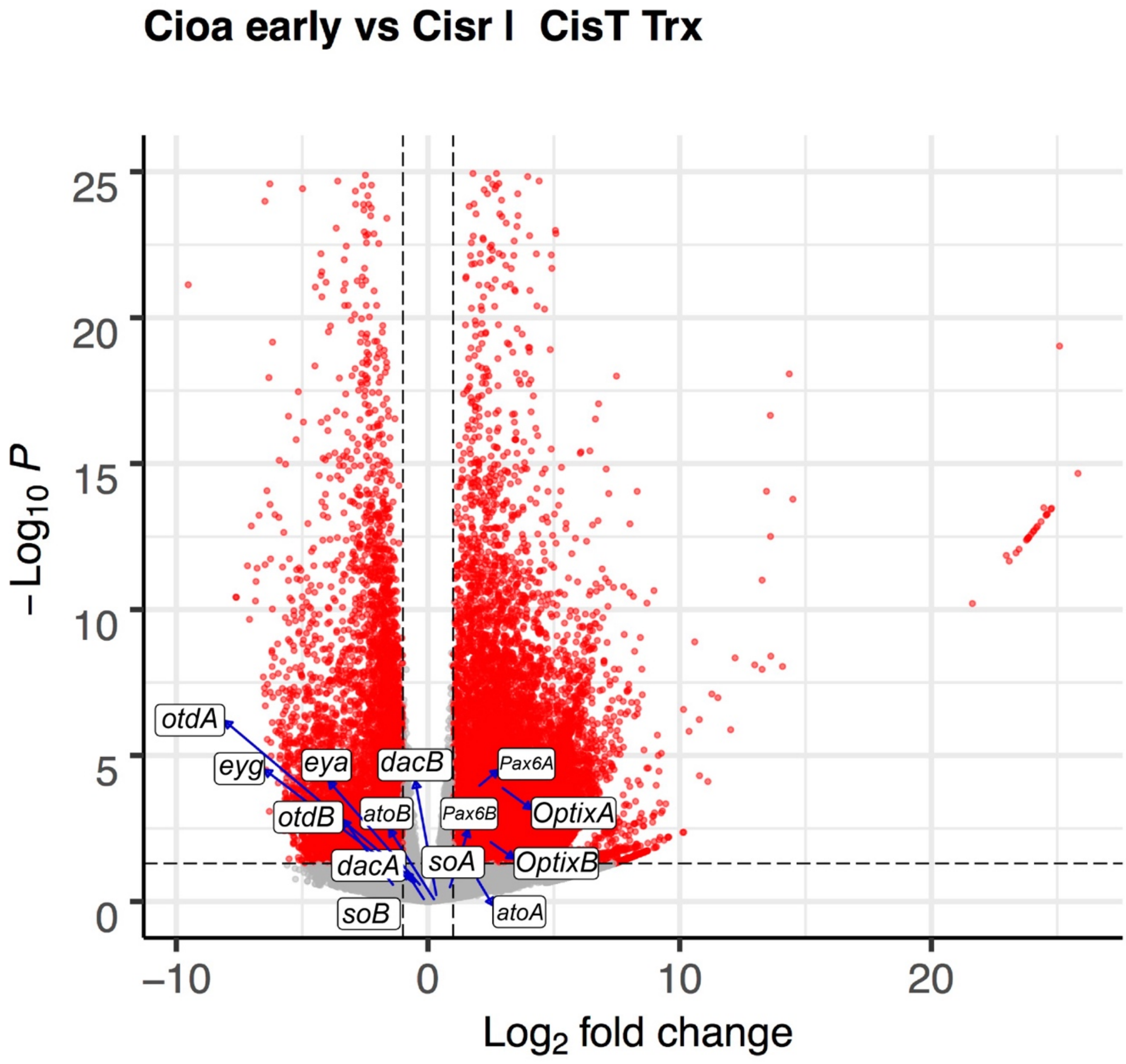


***SI Appendix, Figure S10:*** Comparison 2.1. Volcano plot of $p_{adj}$ values (y-axis) and $log_2$ fold change (x-axis) of all the genes in the analysis differential gene expression comparing reads from early deutembryos of *C. ioanniticus* (normal-eyes) versus early deutembryos of *C. israelensis* (reduced-eyes) mapped onto *C. israelensis* transcriptome. Each gene is represented by a dot. Red dots have $p_{adj}$>0.05. Dashed lines mark $log_2FC$ > [1].

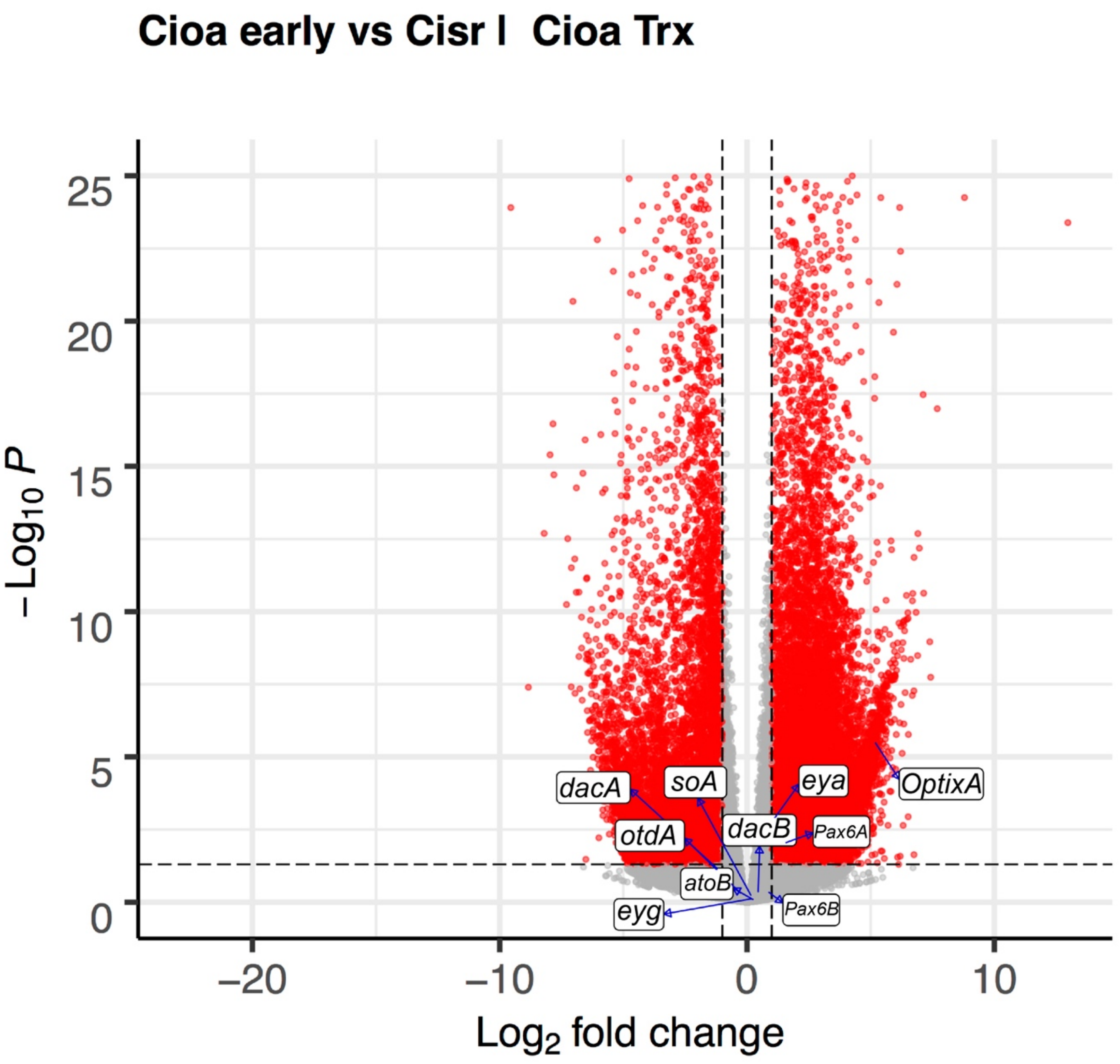


***SI Appendix, Figure S11:*** Comparison 2.2. Volcano plot of $p_{adj}$ values (y-axis) and $log_2$ fold change (x-axis) of all the genes in the analysis differential gene expression comparing reads from early deutembryos of *C. ioanniticus* (normal-eyes) versus early deutembryos of *C. israelensis* (reduced-eyes) mapped onto *C. ioanniticus* transcriptome. Each gene is represented by a dot. Red dots have $p_{adj}$>0.05. Dashed lines mark $log_2FC$ > [1].

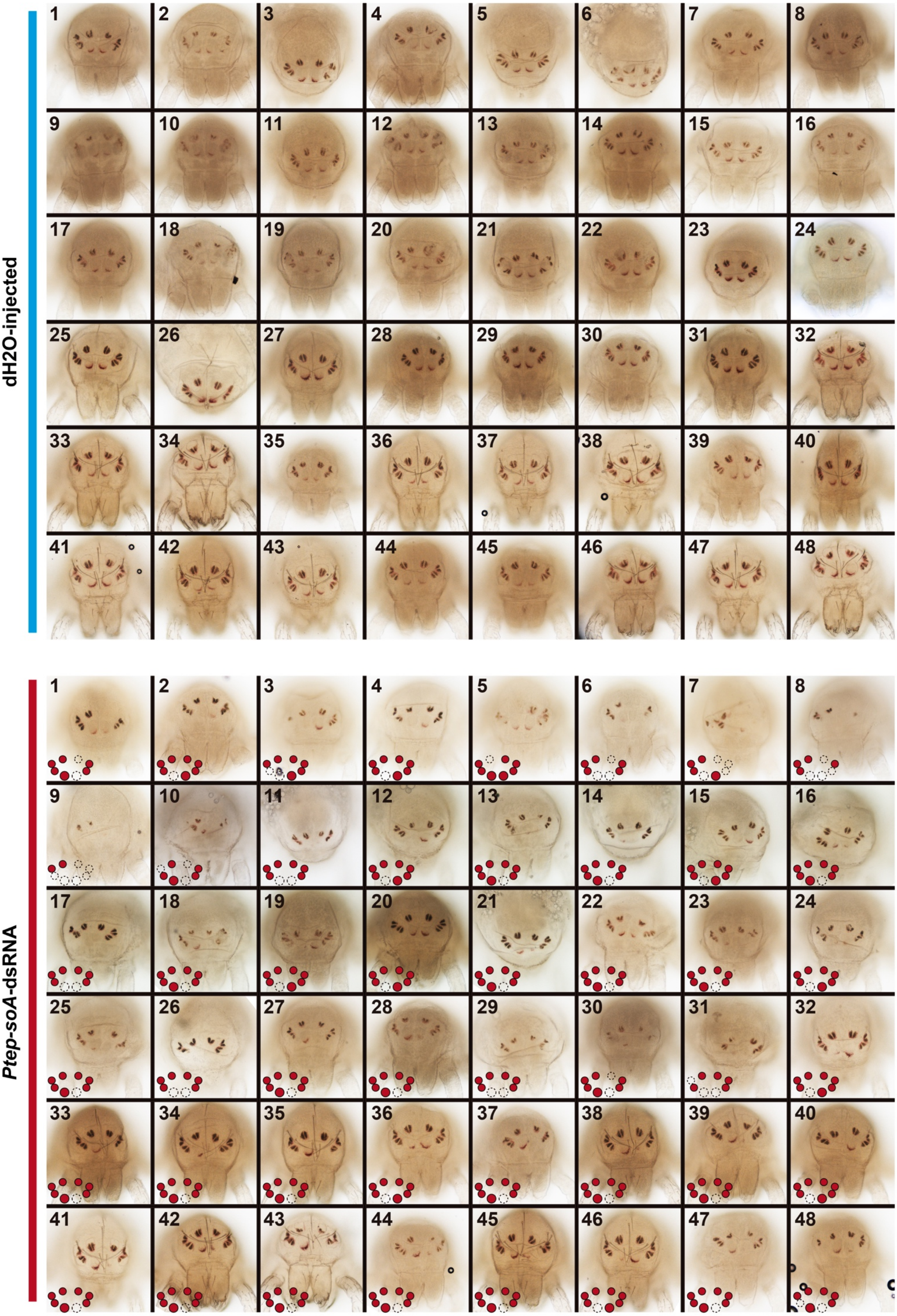
dH2O-injected
Ptep-soA-dsRNA

***SI Appendix, Figure S12:*** Phenotypic spectrum of $dH_2O$-injected (upper panel) and *Ptep-soA*-injected treatments used in the quantification of the effect per eye. High resolution image available at SI Appendix Dataset S3.

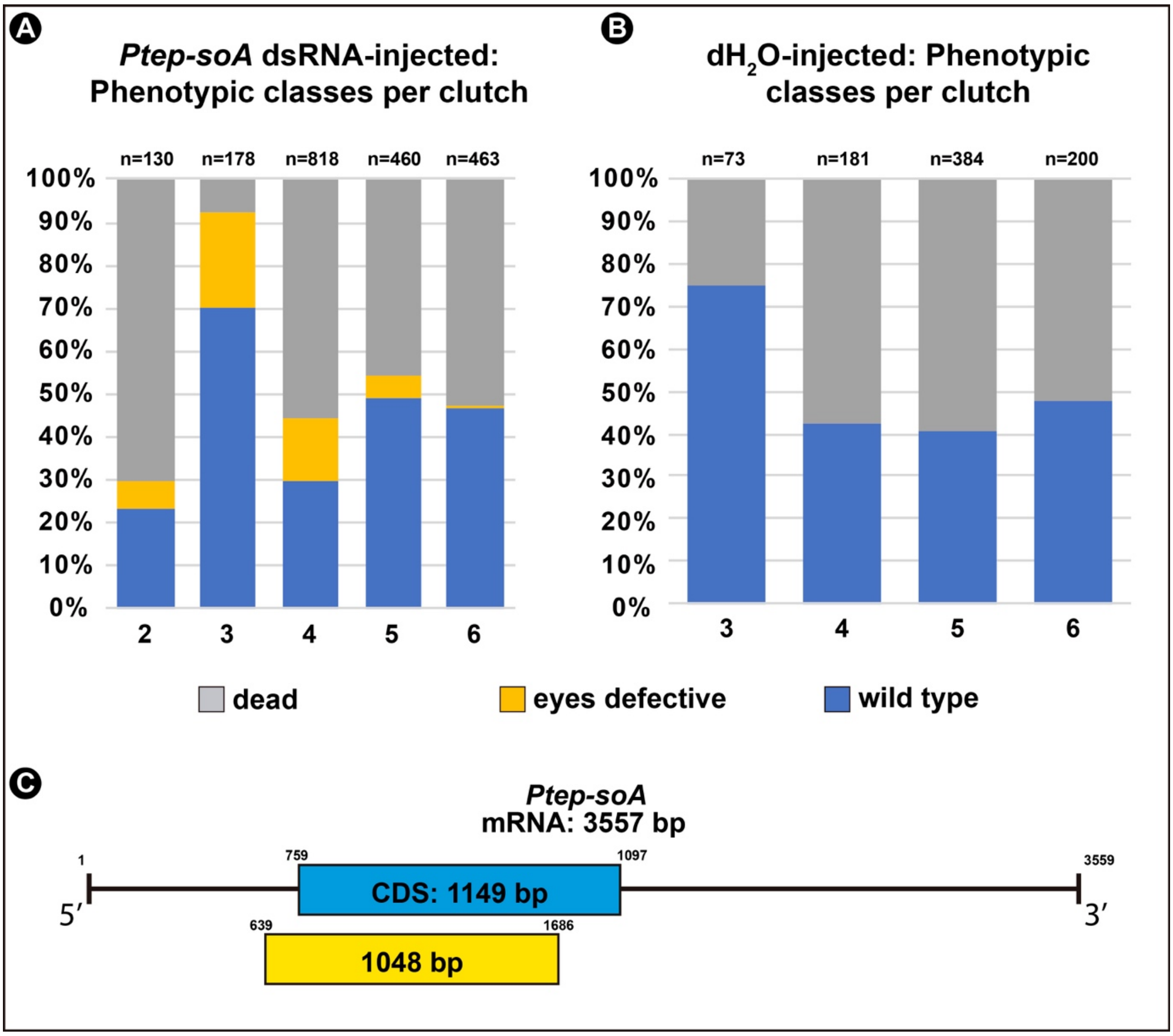


***SI Appendix, Figure S13:*** A. Proportion of individuals in each phenotypic classes per clutch number of the *Ptep-soA* dsRNA-injected treatment. B. Proportions of individuals in each phenotypic classes per clutch number of the $dH_2O$-injected treatment (control). Grey bars: dead; Yellow bar: eyes defective; Blue bar: wild type. C. Schematic representation of *Ptep-soA* transcript, with coding sequence (CDS) highlighted in blue. Yellow box represents the 1048 bp cloned fragment used for probe and dsRNA synthesis.

***SI Appendix, Video S1:*** Time-lapse imaging of a postembryo ~24h after hatching of Parasteatoda tepidariorum from the dH2O-injected treatment (control). Pictures were taken every 30 minutes, in a room at 22ºC. Normal molting time after hatching is ~48h at 26ºC. [downloadable file]

***SI Appendix, Video S2:*** Time-lapse imaging of a postembryo ~24h after hatching of Parasteatoda tepidariorum from the *Ptep-soA*-injected treatment (control). Pictures were taken every 30 minutes, in a room at 22ºC. Normal molting time after hatching is ~48h at 26ºC. [downloadable file]

***SI Appendix, Dataset S1:*** Alignments and sequences of *Charinus* RDGN genes identified in this study.

***SI Appendix, Dataset S2:***

-DESeq2 dataset (*DESeq(dds)*; filtered for $p_{adj}$>0.5) of the DGE analysis of Comparison 1, 2.1 and 2.2 (see "Material and Methods" for explanation)

-*run_ddseq2.r*: custom R script used to run all three analysis

***SI Appendix, Dataset S3:***

-Spreadsheets with the raw counts and sum of counts used to generate the distribution bar plots of the *Ptep-soA* RNAi experiment.

-Primer sequences for the amplified fragments of *Ptep-soA*, *Ptep-otdB* and *Ptep-OptixB*.

-Cloned fragments and amplicon sequences.

-High resolution SI Appendix Figure S12.

***SI Appendix, Table S1:*** Collecting details and sample description for all sequenced samples and respective vouchers.

| **Taxonomy** | **Collecting date** | **ID** | **Locality** | **Longitude** | **Latitude** | **Elevation (m)** | **Preservation** | **Use** | **Description** |
|---|---|---|---|---|---|---|---|---|---|
| *Charinus ioanniticus* | 7/19/18 | ISR021-2 | Hirbet Haruba Cave, deeper chamber | 34.96083 | 31.91328 | 189 | RNAlater | DGE | embryos of female 2 (pre-eye stage); n = 10 |
| *Charinus ioanniticus* | 7/19/18 | ISR021-3 | Hirbet Haruba Cave, deeper chamber | 34.96083 | 31.91328 | 189 | RNAlater | transcriptome assembly; DGE | embryos of female 3 (pre-eye stage); n = 13 |
| *Charinus ioanniticus* | 7/19/18 | ISR021-4 | Hirbet Haruba Cave, deeper chamber | 34.96083 | 31.91328 | 189 | RNAlater | transcriptome assembly; DGE | embryos of female 4 (eye stage); n = 13 |
| *Charinus ioanniticus* | 7/19/18 | ISR021-5 | Hirbet Haruba Cave, deeper chamber | 34.96083 | 31.91328 | 189 | FA/PBST | voucher morphology | embryos of female 2 (pre-eye stage); n = 2 |
| *Charinus ioanniticus* | 7/19/18 | ISR021-6 | Hirbet Haruba Cave, deeper chamber | 34.96083 | 31.91328 | 189 | FA/PBST | voucher morphology | embryos of female 3 (pre-eye stage); n = 2 |
| *Charinus ioanniticus* | 7/19/18 | ISR021-7 | Hirbet Haruba Cave, deeper chamber | 34.96083 | 31.91328 | 189 | FA/PBST | voucher morphology | embryos of female 4 (eye stage); n = 13 |
| *Charinus israelensis* | 7/27/18 | ISR051-4 | Cistern inside Mimlach Cave | 35.44411 | 32.85815 | 139 | RNAlater | transcriptome assembly | embryos of female 2 (deutembryo); n=5 |
| *Charinus israelensis* | 7/27/18 | ISR051-5 | Cistern inside Mimlach Cave | 35.44411 | 32.85815 | 139 | FA/PBST | voucher morphology | embryos of female 2 (deutembryo); n=2 |
| *Charinus israelensis* | 7/27/18 | ISR051-6 | Cistern inside Mimlach Cave | 35.44411 | 32.85815 | 139 | RNAlater | transcriptome assembly; DGE | embryos of female 3 (deutembryo); n=10, |
| *Charinus israelensis* | 7/27/18 | ISR051-7 | Cistern inside Mimlach Cave | 35.44411 | 32.85815 | 139 | FA/PBST | voucher morphology | embryos of female 3 (deutembryo); n=2 |

***SI Appendix, Table S2:*** Summary statistics of transcriptomes of *Charinus ioanniticus* and *Charinus israelesis*.

| **Species** | **Total trinity genes** | **Total trinity transcripts** | **% GC** | **Median contig length** | **Mean contig length** | **Total assembled bases** | **Contig N50 (all transcripts)** | **Median contig length (longest gene)** | **Average contig (longest gene)** | **Total assembled bases (longest)** | **BUSCO** |
|---|---|---|---|---|---|---|---|---|---|---|---|
| *Charinus ioanniticus* | 170848 | 219797 | 39.23 | 332 | 651.88 | 143282365 | 1122 | 305 | 552.92 | 94466089 | C:93.8%[S:88.1% ,D:5.7%],F:3.8%, M:2.4%,n:1066 |
| *Charinus israelensis* | 477982 | 663281 | 38.58 | 323 | 630.61 | 418268343 | 1045 | 297 | 481.28 | 230044656 | C:95.2%[S:88.4% ,D:6.8%],F:2.5%, M:2.3%,n:1066 |
